# Surface-induced tau condensation generates a selective microenvironment around microtubules

**DOI:** 10.64898/2026.08.20.745963

**Authors:** Eva Lanska, Aniruddha Nagarajan, Tereza Humhalova, Valerie Siahaan, Jochen Krattenmacher, Michaela Dujava Zdimalova, Ayoub Belaid, Sinda Khanfir, Lenka Libusova, Carsten Janke, Zdenek Lansky, Marcus Braun, Sandeep Choubey

## Abstract

Tau is a neuron-specific microtubule-associated protein that can self-associate into pathological insoluble aggregates or phase separate into condensates whose (patho)physiological role is debated. Recent studies suggest that intracellular surfaces can locally promote biomolecular condensation, even at low molecular concentrations. While microtubules in neurons provide an extensive tau-interaction surface, their role in tau phase separation remains unclear. Employing theory and in vitro experiments, we here demonstrate that tau forms multilayered condensates on microtubules at physiological concentrations via a prewetting-like transition. Concomitant tau–microtubule and tau–tau interactions explain the experimentally observed cooperative binding of the innermost tau layer directly adsorbed to the microtubule. The formation of this layer is dictated by the spacing of tubulin dimers within the microtubule lattice. Additional tau layers, driven by tau–tau interactions and independent of lattice spacing, are finite in thickness and unstable away from the microtubule surface. While the microtubule-adsorbed tau can selectively restrict proteins from the microtubule surface, the multilayered tau condensates can recruit tau interactors, such as RNA or soluble tubulin, highlighting the distinct roles of the condensate layers. Our results suggest that a prewetting-like transition constitutes a general physical mechanism for organizing liquid-like biomolecular layers of defined composition on charged intracellular surfaces.

**Graphical abstract:** Driven by a pre-wetting-like transition, at concentrations about two orders of magnitude below the critical concentration required for phase separation in solution (which would be required for any possible Plateau-Rayleigh instability), through cooperative binding, tau forms multilayered condensates enveloping microtubules. The condensates consist of an adsorbed layer of slow-turnover tau molecules interacting directly with the microtubule surface (protective against e.g. severing enzymes) and several layers of high-turnover condensed tau sustained by tau-tau interaction only (selectively recruiting e.g. free tubulin).

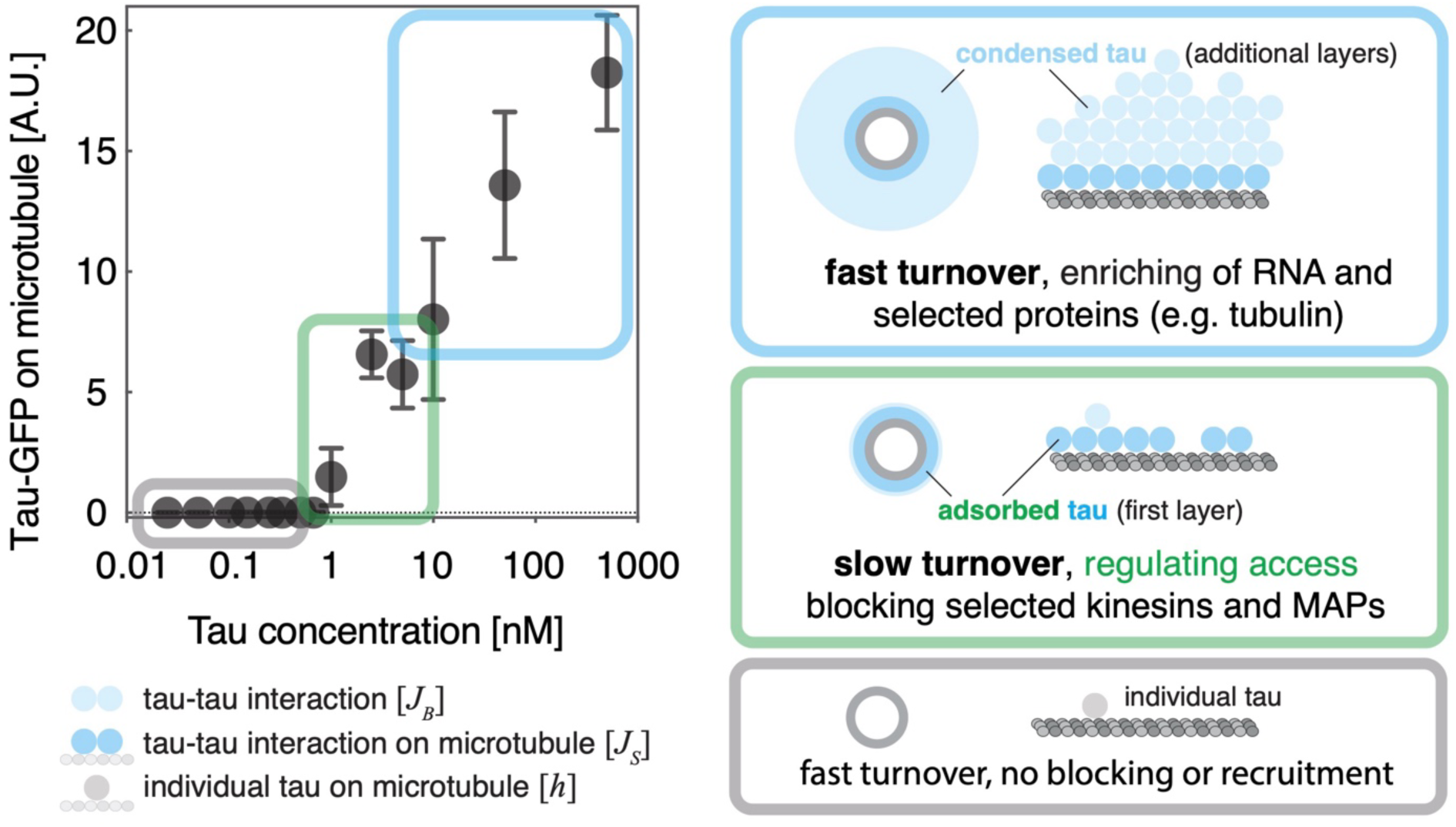

## Introduction

Microtubules are dynamic, cylindrical polymers composed of α- and β-tubulin dimers. They are diverse polymers assembled from heterogeneous tubulin isotypes altered by post-translational modifications^1^. The lattices of mature microtubules are intrinsically plastic, longitudinally compacted or extended, and can exist in distinct conformational states^2,3^. Switching between these conformations can be driven e.g. by binding of specific microtubule-associated proteins^4–8^ or binding of small molecule interactors, like the chemotherapeutic drug taxol^2,9^. Microtubules are essential for diverse cellular functions, such as maintaining cell shape, facilitating intracellular transport, and organizing the mitotic spindle during cell division. In neurons, microtubules act as polarized tracks, enabling long-distance organelle and vesicle transport. Their functions are regulated by a variety of microtubule-associated proteins, many of which are intrinsically disordered^10^, such as the neuron-specific protein tau.

The dysregulation of tau is involved in severe neurodegenerative pathologies, collectively termed tauopathies. The hallmark of tauopathies is the displacement of tau from microtubules and subsequent formation of insoluble tau fibrils via tau-tau interactions. Notably, at micromolar concentrations, tau-tau interactions can also induce liquid-liquid phase separation of tau, leading to the formation of tau droplets, which can recruit specific proteins, including tubulin^11,12^. The roles played by tau liquid-liquid phase separation in physiology and pathology are a subject of ongoing intensive investigation and discussion^13,14^. *In vitro* approaches demonstrated that at picomolar concentrations tau molecules interact with microtubules individually, moving diffusively along the microtubule surface^15^. At low nanomolar concentrations cooperative binding of tau molecules results in the formation of cohesive patches of high tau densities, referred to as ‘islands’, ‘condensates’, or ‘envelopes’ in the literature^6,16,17^. Upon cooperative binding, tau molecules selectively regulate the binding of other microtubule-associated proteins, including molecular motors, to the microtubule surface where kinesin-1 motility is impeded, while the movement of kinesin-8, kinesin-3 and dynein is allowed^16–19^. Moreover, cooperatively bound tau molecules block microtubule-severing enzymes, such as katanin and spastin, thereby protecting microtubules from degradation^6,16,17^. The physical principles governing tau-microtubule association, however, remain elusive.

On attractive surfaces, phase separating proteins can form liquid-like condensates at concentrations below the critical threshold concentration for bulk phase separation in solution. The formation of such surface condensates, driven by a prewetting transition, describes how a dense phase, unstable in solution (in the bulk), can become stabilized at an interface^20–22^. Recent work, extending this concept to biological systems, suggested that prewetting-like transitions may underlie the condensation of proteins on diverse intracellular surfaces, including actin filaments, DNA and lipid membranes^23–25^. Here, using a minimal lattice-gas model in combination with *in vitro* reconstitution experiments, we show that an interplay between tau-tau and tau-microtubule interactions leads to tau condensation on microtubule surfaces, driven by a prewetting-like transition. We show that, as tau concentration increases, tau molecules first assemble into a dense cohesive monolayer, directly adsorbed onto the microtubule lattice, followed by the sequential addition of further tau layers, driving the formation of a multilayered condensate. The formation of the first monolayer, which has been shown to selectively restrict other proteins from the microtubule surface, depends on the surface properties of the microtubule lattice and can be regulated by tubulin post-translational modifications. By contrast, further tau layers can recruit tau-interacting proteins, such as soluble tubulin, thereby locally enabling specific processes, such as enhancing microtubule polymerization through an increase in local tubulin concentration. Employing a dialogue between theory and experiments, our results thus delineate the role of the microtubule surface in the formation of tau condensates, and suggest their functional consequences.

## Results

### Condensation of tau on microtubule lattices is driven by a prewetting-like transition

Tau affinities for microtubules are regulated by the conformation of the tubulin forming the microtubule lattice. The native GDP-bound microtubule lattice, in the absence of stabilizing agents, is predominantly compacted^2,3^, resulting in high affinity cooperative binding of tau^6^. Microtubules stabilized by the non-hydrolyzable GTP analogue GMPCPP are irreversibly extended^2^, resulting in non-cooperative tau binding and much lower tau-microtubule affinity^6^. Microtubules stabilized by the addition of taxol, on the other hand, are reversibly extended^2,3^. As tau binds to taxol-stabilized microtubules it locally compacts the taxol-extended lattice by replacing taxol from the microtubule lattice, resulting in cooperative tau-microtubule interaction albeit with lower affinity compared to the native GDP-lattice^6^.

To study tau-microtubule interactions, we thus considered three microtubule types: (i) compacted GDP-microtubules, (ii) irreversibly extended GMPCPP-stabilized microtubules and (iii) reversibly extended taxol-stabilized microtubules. GDP-microtubules were capped with GMPCPP extensions to prevent depolymerisation following their GTP-driven polymerisation (Methods). The three microtubule types were attached to coverslip surfaces and fluorescently tagged tau was added to follow the kinetics of their interaction using TIRF microscopy (Fig. 1A,B). In line with previous reports^6,16,17^, we observed the formation of high-density tau patches, growing from their edges, which is a hallmark of highly cooperative binding of tau molecules to the microtubule lattice. On GDP-microtubules the high-density tau patches rapidly covered the entire microtubule at 5 pM tau (Fig. 1B left, C left), while no binding of tau was detected on taxol-stabilized and GMPCPP-stabilized microtubules at this concentration (Fig. 1B top, 1D). On taxol-stabilized microtubules, we observed the formation of the characteristic high-density tau patches, growing from their edges, eventually reaching steady-state lengths, at 5 nM tau, three orders of magnitude higher than on GDP-microtubules (Fig. 1B centre, C centre, Movie S1A). At this concentration, the amount of bound tau differed markedly among the three microtubule types (Fig. 1B, E), while at 500 nM tau all three types of microtubules were uniformly covered with tau at similarly high, saturated, densities (Fig. 1B bottom, F). As reported previously, on GMPCPP-stabilized microtubules, at all concentrations tested (5 pM, 5 nM and 500 nM tau), patches were not observed (Fig. 1B right, C right), confirming that tau cooperative binding requires plasticity of the microtubule lattice, namely the reversibility of lattice extension^6^.

**Figure 1.**
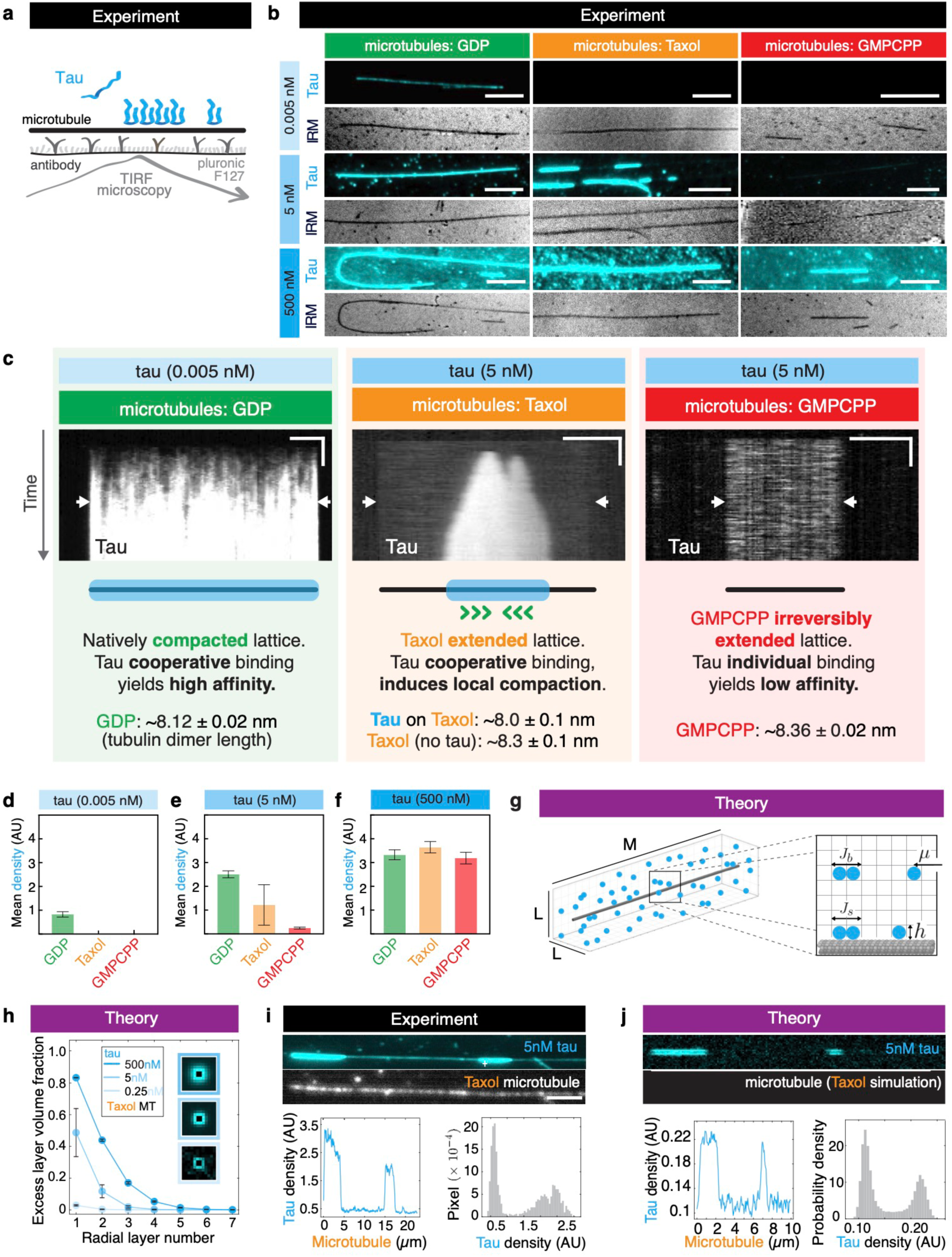
Condensation of tau on microtubule lattices is driven by a prewetting-like transition. **(A)** Schematic of the in vitro experimental setup **(B)** Fluorescence micrographs showing binding of tau–NG (cyan) at different concentrations (0.005, 5, and 500 nM) to distinct microtubule lattices (GDP, taxol-stabilized, and GMPCPP-stabilized). Scale bars, 5 µm. **(C)** Kymographs showing tau assembly on GDP-(0.005 nM tau), taxol-stabilized (5 nM tau), and GMPCPP-stabilized microtubules (5 nM tau). Scale bars: horizontal, 5 µm; vertical, 20 s. Concentrations were chosen to capture initial tau binding dynamics. **(D–F)** Mean intensity of tau on the three microtubule types at 0.005 nM, 5 nM and 500 nM tau concentrations, respectively. **(G)** Schematic of the lattice-gas model of tau interacting with microtubules, parameterized by J_b_, J_s_, h_i_ and μ. **(H)** Simulation results showing the multilayered nature of tau condensates formed on the microtubules. Plot shows the excess tau density (volume fraction of each layer in excess of the bulk volume fraction) at each layer moving radially outward from the microtubule, for 0.25, 5 and 500 nM tau. Inset: simulation snapshots showing end-on view of the microtubule and the tau condensates around it. The concentration values shown are converted from the bulk chemical potential μ (see Supplementary Text for details of the conversion). **(I)** Fluorescence micrograph showing tau binding to a taxol-stabilized microtubule, 4 min after addition of 5 nM tau–NG. Bottom left: fluorescence intensity profile along the microtubule, revealing two populations—high-density tau (condensates) and low-density tau (individually binding tau molecules). Bottom right: histogram of tau signal on the microtubule. Scale bar, 5 µm. **(J)** Simulation results showing condensates formation on a homogeneous surface, demonstrating coexistence of sparse and dense regions on the same microtubule. Bottom left: density profile along the microtubule. Bottom right: histogram of surface tau density from 300 simulations; the bimodal distribution indicates coexistence of two states. Parameters: J_b_ = 0.2 k_B_T, J_s_ = 0.5 k_B_T, h_i_ = 0.0 k_B_T, μ = −2.905 k_B_T. All data in D, E, F and H is presented as mean ± standard deviation (SD).

To gain mechanistic insight into these experimental observations, we employed a lattice-gas model of tau–microtubule interactions. In this model, the microtubule is represented as a cylindrical surface embedded in a three-dimensional cubic lattice, with periodic boundaries (Fig. 1G). The lattice is of size *M* along the microtubule axis, corresponding to the average microtubule length used in the experiments (see Supplementary Text), and size *L* in the transverse directions. Tau molecules occupy the vertices of the lattice, with site occupancy *n_i_* = 1 for an occupied site, *n_i_* = 0 otherwise. Each axial position contains 12 radially arranged tau binding sites, mimicking the protofilaments of a microtubule. One binding site is taken to be approximately 30 nm in size (refer to Supplementary Text). The Hamiltonian dictating tau binding to the microtubule surface is given by

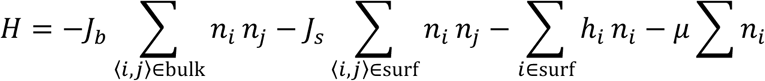

The first term, characterized by the coupling parameter *J_b_*, accounts for the nearest-neighbour interactions between tau molecules in solution (also referred to as the bulk), reflecting the multivalent nature of tau and its propensity to form phase separated condensates^11^. The second term, parameterized by *J_s_*, captures interactions between neighboring tau molecules bound to the microtubule surface, accounting for tau cooperativity^16^. The third term represents binding of tau to the microtubule, with site-dependent affinity *h_i_* at binding site *i*. Note that the second and third terms contribute only at sites adjacent to the microtubule surface. The final term, with bulk chemical potential *μ,* sets the concentration of tau in the solution. In the dilute limit, *μ* can be related to the tau concentration in the experiments, enabling quantitative comparison with the data (refer to Supplementary Text and Supplementary Fig. S1 for details).

We modeled tau binding behaviour to the three experimentally studied microtubule lattices through appropriate choices of *J_s_* and *h_i_*, initially focussing on taxol-stabilized microtubules. We first considered that the microtubule surface is homogeneous for tau binding, i.e., we assigned the same value of *h_i_* to every binding site along the microtubule. We refer to this as the “homogeneous-surface model” (Supplementary Fig. S2). To characterize the equilibrium properties of this model, we performed Monte Carlo simulations at tau concentrations below the threshold for phase separation in solution, reported to be around 5 μM to 10 μM^13,26^. At very low concentrations (< 1 nM) tau molecules get sparsely and individually adsorbed onto the microtubule surface (Fig. 1H, inset bottom). As the tau concentration is increased, cooperative binding of tau on the microtubule surface leads to the formation of cohesive patches or condensates (Movie S1). At these intermediate concentrations (∼ 1-10 nM), the model predicts coexistence of high and low-density regions of tau along the microtubule lattice (Fig. 1H inset middle, Fig. 1J). This coexistence is reflected in the bimodal distribution of tau densities, consistent with experimental findings (Fig. 1I, J). Although the tau population on the microtubule is dominated by molecules directly adsorbed to the surface, the model predicts that tau–tau interactions also promote the formation of less stable secondary layers above the adsorbed layer which already occurs in tau condensates at intermediate concentrations. Subsequent increase in tau concentration (> 100 nM) predicts the formation of multilayered condensates (Fig. 1H, inset top) formed via a prewetting-like transition. Repeating the simulations for GDP-and GMPCPP-stabilized microtubules provided a mechanistic explanation for how microtubules with distinct surface properties exhibit comparable tau densities at high protein concentrations (Fig. 1B bottom, F). The initial adsorption of tau is governed by its affinity for the underlying microtubule surface; low-affinity surfaces, such as GMPCPP-stabilized microtubules, exhibit reduced tau binding at low concentrations. However, as the concentration increases, more tau molecules begin to bind the microtubule surface. Once the first layer of tau forms, additional layers of tau can assemble through tau–tau interactions, independent of microtubule surface chemistry, resulting in similar overall intensities at saturating concentrations (Supplementary Fig. S2F). Throughout this manuscript, we use the term “adsorbed tau” to refer to the initial layer of tau molecules that directly binds the microtubule lattice, and “condensed tau” for the subsequent layer(s) of tau that are interacting with the adsorbed layer through tau-tau interactions. Combined, these results provide a unified theoretical framework for tau-microtubule interaction and predict the existence of multilayered tau condensates on the microtubule lattice formed via prewetting-like transition.

### Microtubule surface heterogeneity dictates spatial patterns of tau condensation

To develop a bottom-up understanding of the principles governing tau condensation on microtubules, we first investigated how the interplay between tau–microtubule and tau–tau interactions give rise to the observed spatial patterns of condensate nucleation and formation (Fig. 2A). To experimentally access the question of localized nucleation of the condensates, we employed taxol-stabilized microtubules, which readily enable visualization of the cooperatively-bound tau as high-density tau patches interspersed by low-density tau regions, where tau molecules bind independently (Fig. 2A). Complete washout of tau from the channel after establishing tau condensates on the microtubules enabled us to repeatedly access the positions of condensate nucleation experimentally on the same sets of microtubules. Experimental kymographs reveal that after tau flush out (Fig. 2A lower kymograph), tau condensates have a strong tendency to form at similar positions along the microtubule (Fig. 2B – left, Movie S2A). Our homogeneous-surface model fails to explain these experimental observations; the model predicts that tau condensates nucleate stochastically at arbitrary positions along the microtubule. Furthermore, in contrast to the experimental observations, the model predicts that on homogeneous microtubules, tau condensates do not remain anchored to their initial nucleation sites, but instead can continuously translocate along the microtubule lattice (Fig. 2B centre, Supplementary Fig. S2A).

**Figure 2.**
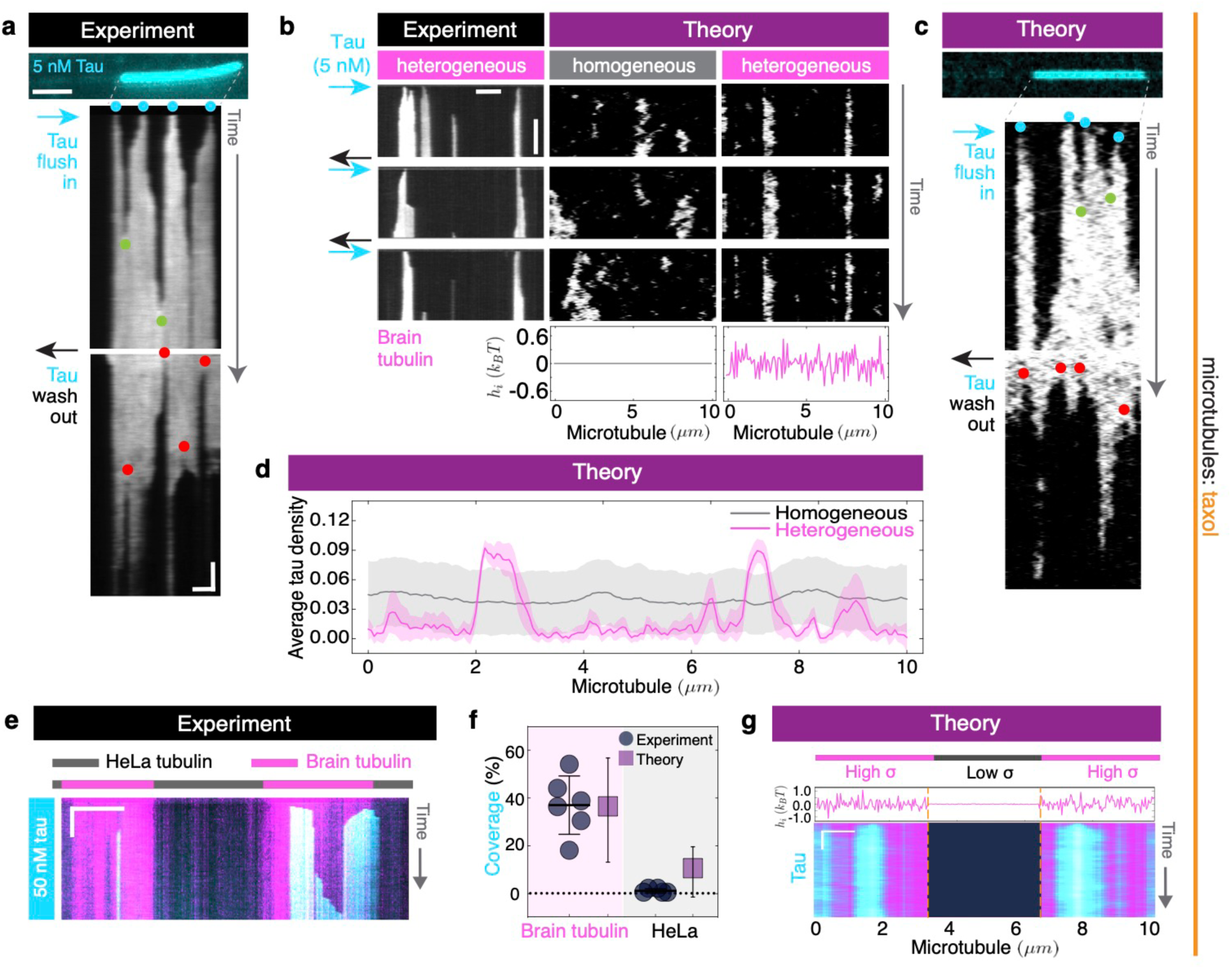
Microtubule surface heterogeneity dictates spatial patterns of tau condensation. **(A)** Multichannel fluorescence micrograph showing tau–mNG (cyan) signal on a taxol-stabilized microtubule. Kymograph (middle) shows condensate formation and growth following addition of 5 nM tau-mNG. Initially, the microtubule is sparsely coated; over time, high-density tau regions emerge. Cyan dots indicate the tau condensate nucleation site, and the green dots show fusion sites of adjacent condensates. A second kymograph (bottom) shows condensate disassembly after removal of tau from solution; the red dots indicate the fission events. Scale bars: horizontal, 2 µm; vertical, 1 min. **(B)** Left: kymographs showing tau binding over three cycles of tau addition (5nM) and removal. Tau condensates consistently assemble at the same regions, indicating spatially selective binding. Scale bars: horizontal, 5 µm; vertical, 5 min. Centre: kymographs from independent simulations on a homogeneous surface (h_i_ = 0); condensates form at variable locations and are not spatially pinned. Right: independent simulations on a heterogeneous surface (h_i_ ∼ N(0.0, 0.2)); condensates form at fixed locations and remain pinned. Parameters: J_b_ = 0.2 k_B_T, J_s_ = 0.5 k_B_T, μ = −2.91 k_B_T. **(C)** Simulation kymographs showing condensate assembly (top) and disassembly (bottom) on a heterogeneous surface. Parameters: J_b_= 0.2 k_B_T, J_s_ = 0.5 k_B_T, μ = −2.91 k_B_T, and h_i_ ∼ N(0.0, 0.2). **(D)** Tau density profile along the microtubule, averaged over 100 simulations (parameters as in B). **(E)** Kymographs illustrating tau binding to the two tubulin types within the annealed microtubule: porcine brain tubulin (magenta-labelled) and HeLa cell tubulin (unlabelled), at 50 nM tau. Condensate formation is strongly reduced on HeLa microtubule sections compared to porcine brain sections. Scale bars: horizontal, 5 µm; vertical, 2 min. **(F)** Quantification of the percentage of microtubule length covered by tau condensates in (E) and (G), showing strong agreement between experiment and simulation. Simulation of an analogous heterogeneous microtubule shows preferential condensate formation on the more heterogeneous regions. In dot plot (experiment), each dot represents one analyzed field of view (containing at least seven annealed microtubules). Data is presented as the mean (line) ± SD (error bars). **(G)** Simulation of an analogous heterogeneous microtubule shows preferential formation on the more heterogeneous regions. Parameters: J_b_ = 0.2 k_B_T, J_s_ = 0.5 k_B_T, μ = −2.935 k_B_T and h_i_ ∼ N(0.0, 0.01) in the central region and h_i_ ∼ N(0.0, 0.2) in the flanking regions (averaged over 50 simulations).

To account for these experimental observations, we extended the model to include spatial heterogeneity along the microtubule surface, potentially arising from variations in tubulin isotypes and post-translational modifications (PTMs)^27,28^, or variations in lattice spacing between tubulin dimers^29^. In this “heterogeneous-surface model”, site-dependent affinity *h_i_* at microtubule binding site *i* is randomly sampled from a normal distribution (see Supplementary Text and Fig. S3 for details). In contrast to the homogeneous-surface model, the heterogeneous-surface model predicts the formation of non-moving, spatially localized condensates, in agreement with our experimental findings (Fig. 2B right, C). Additionally, similar to the experiments, repeated spatial patterns of condensation were conserved across independent simulation runs (Fig. 2B – right, D, Movie S2). Our results so far suggest that the specificity of tau condensates for certain positions along the microtubule is intrinsic to the microtubule lattice itself. Interestingly, the simulations show that an uncorrelated random mixture of tau-microtubules affinities is sufficient to explain the emergence of preferred regions of tau condensate nucleation and growth along the microtubule surface. This suggests that variations in tubulin isotypes and PTMs, potentially in combination with differences in lattice spacing, can explain the spatial specificity of the tau condensates. We interpret this to arise due to the cooperative nature of the tau-microtubule interaction. Regions which, stochastically, contain higher density of tubulin dimers with higher affinity for tau, will locally attract more tau molecules to the microtubule lattice and will thus, through locally increasing probability of tau-tau interactions, present favorable spots for condensate nucleation (refer to Supplementary Fig. S4 for further comparisons to experiments).

To systematically test this hypothesis and shed light on the mechanistic origin of microtubule heterogeneity for tau binding, we performed experiments with microtubules containing defined inhomogeneities in the content of tubulin variants along the microtubule lattice. To this end, we generated hybrid microtubules by annealing fragments of microtubules polymerized from porcine brain tubulin, which have high levels of PTMs, and microtubules polymerized from tubulin isolated from HeLa cells, which have low levels of PTMs and different tubulin isotype distribution. We found that tau condensates form predominantly on the porcine brain microtubule sections (Fig. 2E, F), confirming that differences in types of tubulin can affect the localized nucleation of the tau condensates. Predictions from the model, using an analogous construction of microtubule, corroborated these findings (Fig. 2F, G), further highlighting the role of surface heterogeneity.

In summary, a dialogue between our theory and experiments indicates that heterogeneity in tau affinities along the microtubule surface, likely mediated by tubulin PTMs and isotypes, dictates the spatial pattern of tau condensate formation. Remarkably, our results show that even a random arrangement of these tubulin variants and PTMs is sufficient for the emergence of tau condensates at specific positions along the microtubule.

### Condensates comprise two kinetically distinct tau populations

Our model predicts that the formation of multilayered tau condensates using tau concentrations above threshold concentrations depends on the type of microtubule lattice (GDP-, GMPCPPP-stabilized or taxol-stabilized). Specifically, for taxol-stabilized microtubules, the model predicts that the threshold concentration for surface condensation is at around 1 nM tau (Supplementary Fig.S3B). The first layer of tau adsorbs onto the microtubule through the combined effects of tau-tau cooperativity and tau-microtubule binding affinity, determined by *J_s_* and *h_i_*, respectively. As this adsorbed layer becomes saturated, additional layers of tau are recruited, governed by tau-tau interactions, as characterized by the interaction parameter *J_b_* (Fig. 3A). The model predicts distinct timescales of dissociation for the adsorbed and condensed tau. More specifically, when the combined effect of *J_s_* and *h_i_* is significantly stronger than *J_b_*, the model predicts that upon depleting tau from the solution, the condensed layers will disassemble rapidly, followed by a slower decay of the adsorbed layer (Fig. 3B, Supplementary Fig. S5A). To test this model prediction, we reanalyzed previously published data^16^ from experiments in which we considered taxol-stabilized microtubules with pre-formed tau condensates, then removed tau from the solution, and monitored the change in tau intensity on the microtubule. Strikingly, we observed that tau density in condensates decays with two characteristic timescales: within a couple of seconds, the intensity dropped significantly, revealing a fast-turnover population within the condensates. This fast drop was followed by a much slower decay on the order of tens of seconds, during which the condensate disassembled, revealing the second tau population exhibiting slow turnover (Fig. 3C, Supplementary Fig. S5B).

**Figure 3.**
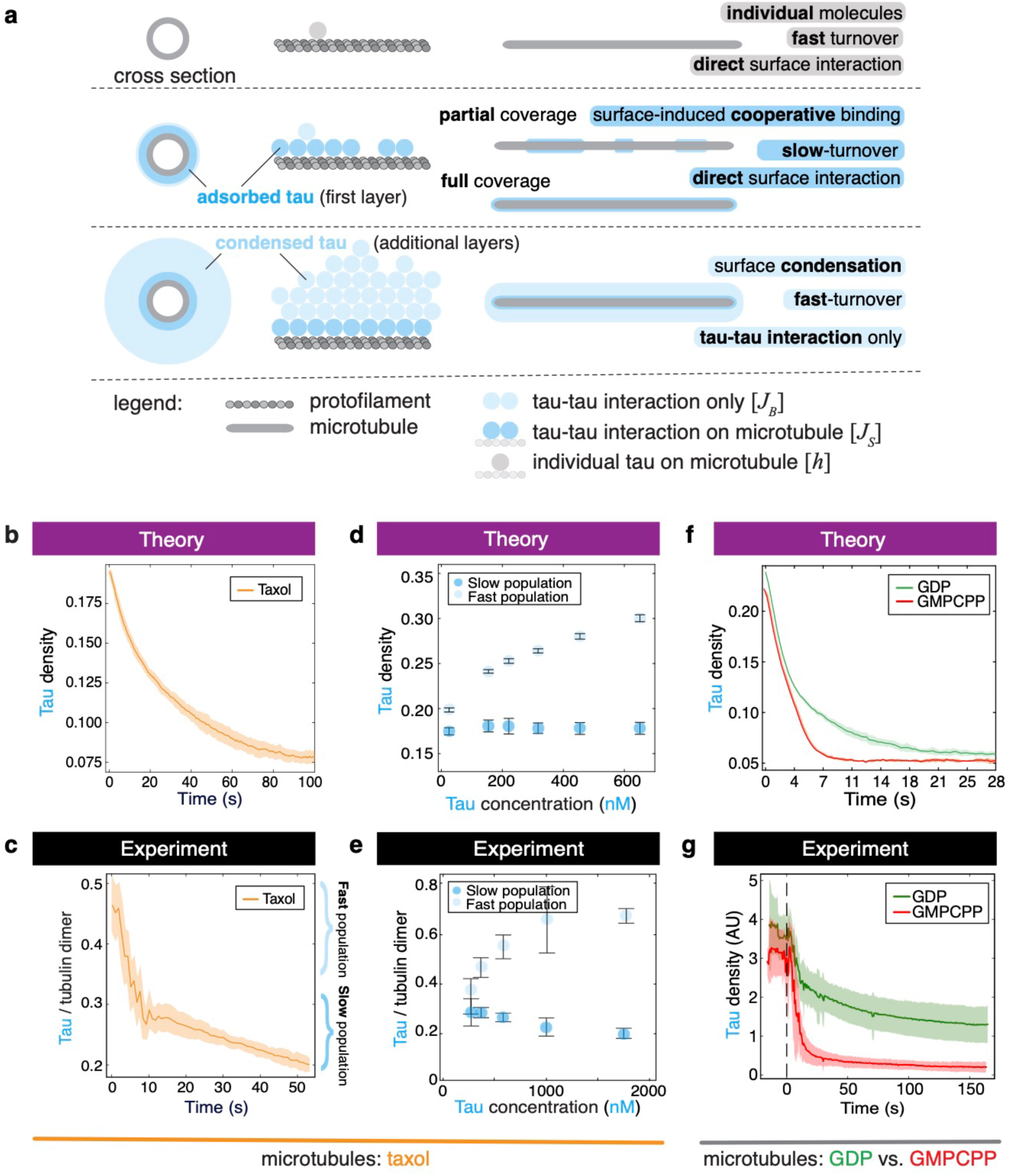
Condensates comprise two kinetically distinct tau populations. **(A)** Schematic illustrating distinct modes of tau binding to the microtubule. **(B)** Simulation of tau density during disassembly, revealing fast- and slow-turnover populations identified by a double-exponential fit (Supplementary Fig. S5A). **(C)** Experimental data showing time-traces of tau-meGFP density during tau washout, showing fast- and slow-turnover components within tau condensates (160 nM tau). Analysis was performed on 9 condensate regions across 3 independent experiments. **(D)** Simulation results showing tau density on the microtubule following the disassembly kinetics at various concentrations. Fast population corresponds to the tau density at the initial stage, before the washout, and slow population corresponds to the density when the timescales switchover happens or when the condensate begins to disassemble from the sides (J_b_ = 0.2 k_B_T, J_s_ = 0.5 k_B_T, h_i_ ∼ N (0.0, 0.2), μ related to concentration using fitting parameters (see Supplementary Text), averaged over 30 simulations. **(E)** Tau densities measured within condensates before and after tau washout at various concentrations. Two populations of tau can again be identified: fast- and slow-turnover tau. The fast-turnover tau population increased, while the slow-turnover tau population remained similar. Analysis was performed on 6 condensate regions across 3 independent experiments. **(F)** Model prediction for condensate disassembly when comparing microtubules with different surface properties, where the condensate is allowed to first form with 500 nM tau concentration. GDP-microtubules are assigned h_i_∼Normal (0.3, 0.2) and GMPCPP-stabilized microtubules have the affinities as h_i_∼Normal (−0.1, 0.2). The model predicts that the fast timescale should be the same in both cases. **(G)** Experimental measurements of tau (500 nM) disassembly kinetics on GDP- and GMPCPP-stabilized microtubules, confirming the model prediction. N=16 GDP- and 19 GMPCPP-stabilized microtubules across 3 independent experiments. All data from B, C, F, G is presented as mean (line) ± SD (shaded areas) and D, E is presented as mean (dot) ± SD (error bars).

Moreover, the model predicts how these two populations would change with increasing tau concentration. Once the adsorbed layer (slow-turnover population) becomes saturated, further increase in tau concentration only leads to the formation of subsequent condensed layers (fast-turnover population). Therefore, the model predicts that the fraction of tau molecules in the slow-turnover population will remain constant with increasing tau concentration, and the fraction of fast-turnover tau will increase with increasing tau concentrations (Fig. 3D). To test this model prediction, we repeated the wash-out experiments at different tau concentrations and quantified the relative magnitude of the two populations in tau condensates. We found that the slow-turnover population remained approximately constant, whereas the fast-turnover population increased with tau concentration and saturated in the micromolar range (Fig. 3E), in close agreement with the model predictions. Monitoring tau turnover on the microtubule outside of the condensates in a similar way showed only one tau population with fast turnover on the order of seconds (Supplementary Fig. S5C, D). This experimental observation is consistent with the theoretical prediction that the regions outside the condensates only consist of a weakly bound layer of tau. The model also predicts, and the experiments confirm, that tau condensates stop growing in thickness at sufficiently high tau concentration as evidenced by the saturation of tau density on the microtubule at elevated tau concentrations (Fig. 3D, E). As shown in previous studies of wetting on cylindrical surfaces^30,31^, this saturation arises due to the curvature of the microtubule, which can be approximated as a thin cylinder. As the condensed layer grows thicker, the interfacial area between the dense and dilute phases increases, leading to an increasing free energy cost. Consequently, further growth of the condensed layer becomes thermodynamically unfavourable, resulting in the observed saturation. Consistently, as we titrated tau to microtubules over a wide range of concentrations and measured the tau signal on the microtubule in equilibrium (Supplementary Fig. S6A), we obtained a complex tau-microtubule binding curve that could not be fitted to a single Hill function, suggesting multiple binding regimes at different tau concentrations. In the low and intermediate concentration regime, the binding exhibited highly cooperative behaviour (Supplementary Fig. S6B, inset), consistent with an adsorbed layer that depends on tau-tau as well as tau-microtubule interactions, while in the high concentration regime, the binding could be well approximated by a non-cooperative Michaelis-Menten behaviour, consistent with the addition of multiple condensed tau layers. A good fit over the entire concentration range was obtained using the heterogeneous model, further substantiating the validity of our theory (Supplementary Fig. S6).

To further probe the multilayered nature of tau condensates, we again turn to the model, which predicts that varying the microtubule surface properties (via *J_s_* and *h_i_*) should not affect the condensed tau, which is the fast-turnover tau population governed by *J_b_* and *μ* (Fig. 3F, Supplementary Fig. S5E). Consistent with this model prediction, experiments performed on GDP- and GMPCPP-stabilized microtubules, with tau condensates formed at 500 nM tau, exhibit identical disassembly timescales for the fast-turnover tau population. This is despite the markedly different microtubule surface properties, which, in the case of GDP-lattices, support the formation of slow-turnover, cooperative, tau population, and in the case of GMPCPP-lattice does not (Fig. 3G, Supplementary Fig. S5F). Conversely, the model predicts that adding different tau concentrations to the same microtubule type does not affect the disassembly timescale of the slow population, which corresponds to the layer adsorbed directly to the microtubule. Accordingly, the slow disassembly timescale is expected to be independent of tau concentration for a given microtubule, as shown in Fig. S5G. This prediction is also corroborated by experiments measuring the disassembly timescales on GDP-microtubules at 10 nM, which should result in the formation of tau condensates formed mainly of adsorbed tau, and 500 nM, where tau condensates should contain a large degree of condensed tau (Supplementary Fig. S5H).

In summary, our model predicts, and our experiments confirm, the existence of two kinetically distinct populations of tau in the condensates. The two populations arise as a consequence of the multilayered nature of the tau condensates, with the slow-turnover tau molecules directly adsorbed onto the microtubule surface, and the fast-turnover tau population corresponding to molecules condensed onto the adsorbed layer at elevated tau concentrations.

### Condensed tau recruits tau interactors to the microtubule surface

A combination of our theory and experiments suggests that, within a physiological range of concentrations^32^, tau forms multilayered condensates on microtubules. We thus wondered whether, analogous to tau droplets formed by liquid-liquid phase separation in solution^11^, these surface-induced tau condensates could recruit tau interactors, such as tubulin. To test this hypothesis, we attached GDP-microtubules to the coverslip surface and probed their interaction with soluble fluorescently labeled tubulin. In the absence of tau, we did not observe any significant binding of soluble tubulin to the microtubule surface (Fig. 4A left, B). Neither was tubulin recruited to microtubules covered by tau condensates formed at 5 nM tau, which contain predominantly the microtubule-adsorbed, slow-turnover tau (Fig. 4A centre, B). Changes of soluble tubulin concentrations over an order of magnitude did not change the outcome of these experiments (Fig. S7A). Strikingly, adding soluble tubulin to microtubules covered by tau condensates formed at 500 nM tau, which also contain the condensed or fast-turnover tau population, led to pronounced recruitment of soluble tubulin to the tau-coated microtubules (Fig. 4A right, B). The amount of tubulin recruited was depended on the concentration of tubulin in solution (Supplementary Fig. S7A). The same effect was observed with taxol- and GMPCPP-stabilized microtubules (Supplementary Fig. S7B, C) suggesting that the fast turnover tau, which is present on taxol- and GMPCPP-stabilized microtubules at elevated tau concentrations, is responsible for this effect (Movie S3).

**Figure 4.**
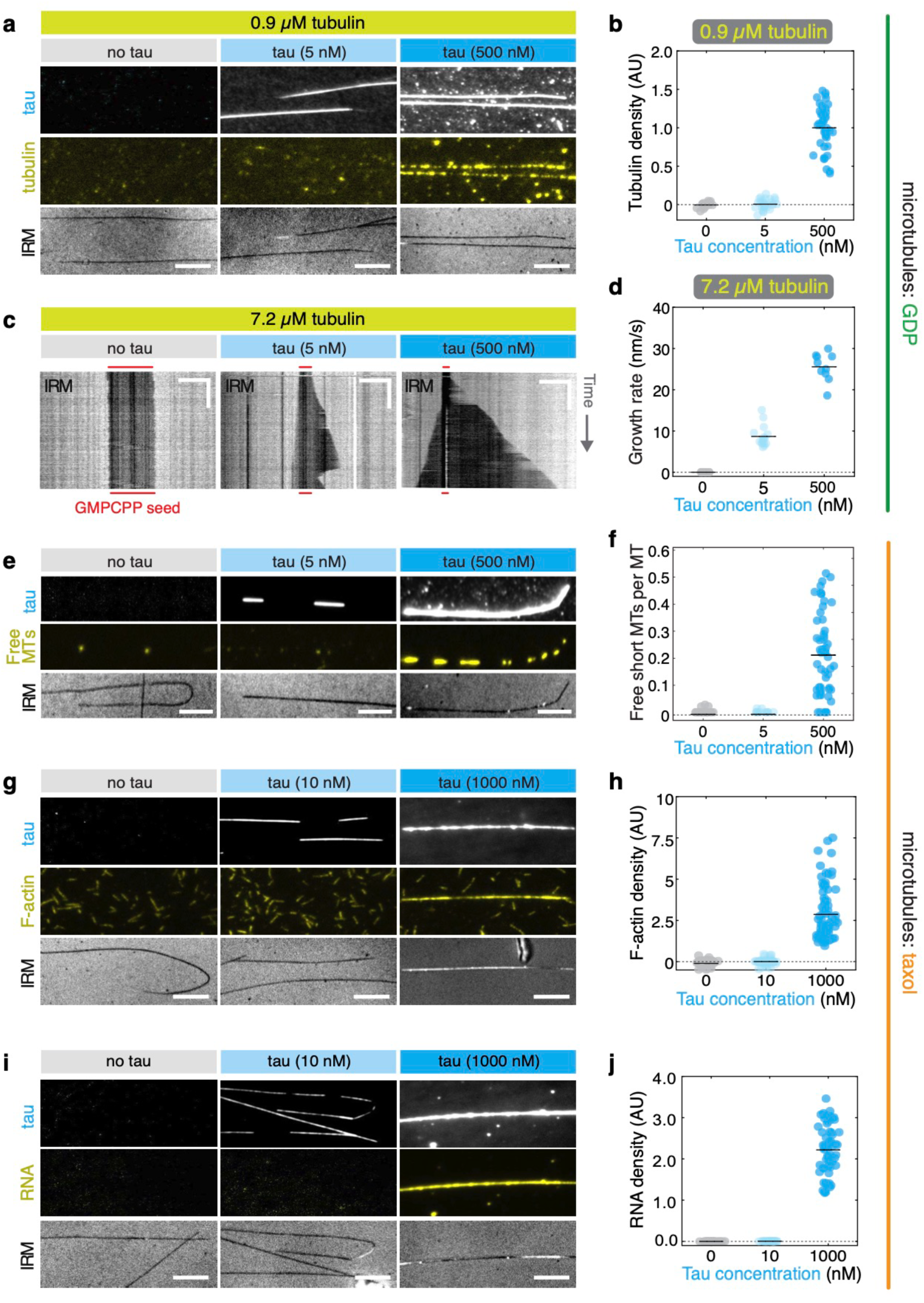
Condensed tau recruits tau interactors to the microtubule surface. (**A)** Fluorescence snapshots showing free tubulin (Hilyte647-labeled) on GDP-microtubules in the presence of 0, 5, or 500 nM tau–mNG. Tau was first added to unlabeled GDP-microtubules, followed by addition of 90 nM Hilyte 647-labeled tubulin in presence of tau. Snapshots correspond to the final frame of a 10 min acquisition. Scale bar, 5 µm. **(B)** Quantification of tubulin fluorescence intensity on microtubules for conditions in (A). N=17–45 microtubules per condition across 3 independent experiments. **(C)** Kymographs showing microtubule growth from GMPCPP seeds after addition of 7.2 µM tubulin in the presence of 0, 5, or 500 nM tau–mNG. Scale bars: 5 µm (horizontal), 3 min (vertical). **(D)** Microtubule growth rates for conditions in (C). N=12–18 microtubules per condition across 3 independent experiments. **(E)** Fluorescence snapshots showing short, Hilyte647–labeled microtubules on immobilized taxol-stabilized microtubules in the presence of 0, 5, or 500 nM tau–mNG. Tau was added first, followed by labeled microtubules in presence of tau. Snapshots correspond to the final frame of a 10 min acquisition. Scale bars, 5 µm. **(F)** Quantification of free short Hilyte647-labeled microtubules on immobilized long microtubules − conditions in (E). N=16–57 microtubules per condition across 2 independent experiments. **(G)** Fluorescence snapshots showing F-actin (phalloidin-stabilized, rhodamine labeled) on taxol-stabilized microtubules in the presence of 0, 10, or 1000 nM tau–mNG. Tau was added first, followed by F-actin in presence of tau. Snapshots correspond to the final frame of a 15 min acquisition. Scale bars, 5 µm. **(H)** Quantification of F-actin fluorescence intensity on microtubules for conditions in (G). N=69–79 microtubules per condition across 3 independent experiments. **(I)** Fluorescence snapshots showing RNA-oligonucleotides localization on taxol-stabilized microtubules in the presence of 0, 10, or 1000 nM tau–mNG. Tau was added first, followed by RNA in presence of tau. Snapshots correspond to the final frame of a 15 min acquisition. Scale bars, 5 µm. **(J)** Quantification of RNA fluorescence intensity on microtubules for conditions in (I). N=32–63 microtubules per condition across 3 independent experiments. In all plots in B, D, F, H, J,, each dot represents one microtubule and the horizontal line denotes the mean.

We next wondered if tubulin partitioning into the tau condensates may have any functional consequences. To address this, we performed *in vitro* polymerization assays using GMPCPP-stabilized microtubule seeds and soluble tubulin. In the absence of tau, no microtubule growth was observed at 7.2 µM soluble tubulin. In contrast, in the presence of tau (5 nM and 500 nM), microtubule elongation was detected under the same conditions. In the presence of 500 nM tau, microtubule growth rates were approximately threefold higher than at 5 nM tau, and catastrophe events were completely abolished (no catastrophes observed at 500 nM tau) (Fig. 4C, D; Supplementary Fig. S7F, Movie S4). A similar trend was observed across all tested tubulin concentrations (Supplementary Fig. S7D, E, Movies S5, 6). We attribute the pronounced effect at 500 nM tau to local enrichment of soluble tubulin at the microtubule surface. At low tau concentration (5 nM), where no detectable tubulin recruitment was observed, tau nevertheless promotes microtubule growth. This suggests that tau can enhance microtubule polymerization through a mechanism that is independent of tubulin partitioning. Nevertheless, at higher tau concentrations tubulin recruitment by the condensed tau layers further enhances microtubule polymerisation.

To test the effect of tau condensation on microtubule bundling, we attached long biotinylated taxol-stabilized microtubules to the coverslip surface via anti-biotin antibodies. After addition of short non-biotinylated microtubules to the solution, we did not observe any microtubule-microtubule interaction neither in the absence of tau nor in the presence of tau condensates formed at low tau concentration (5 nM) where tau condensates are mainly composed of low-turnover adsorbed tau (Fig. 4E, F). However, repeating the experiment with tau condensates formed at 500 nM tau, composed of low- and high-turnover tau, we observed robust formation of microtubule bundles (Fig. 4E, F, Movie S7). Encouraged by this finding, and knowing that tau also interacts with actin^33,34^, we next tested the binding of stabilized actin filaments to our tau-covered microtubules and observed robust binding of actin filaments, again only when tau condensates contained high-turnover condensed tau (Fig. 4G, H, Movie S8). Finally, since phase separated tau is known to recruit RNA^35, 36^, we tested whether fluorescently tagged polyA-RNA can be recruited to the tau condensates and found that again only high-turnover tau-containing condensates recruit RNA to the surface of microtubules (Fig. 4I, J, Movie S9). Taken together, these experiments and theory demonstrate that tau condensates comprise two distinct tau populations with different functional roles. A low-turnover, microtubule-adsorbed tau layer forms the stable core of the condensate, whereas the more dynamic, high-turnover outer layers mediate the recruitment of many tau interactors, including soluble tubulin and neighbouring microtubules and actin filaments. Through this selective enrichment of interacting components, tau condensates can generate localized microenvironments with distinct functional properties, such as increased local tubulin concentrations that promote enhanced microtubule growth.

## Discussion

Although tau has been extensively studied for decades, our understanding of tau physiology and of the processes leading to tau-related pathologies is still limited. Recent results suggest that tau-tau interactions, which have been implicated in the formation of pathological tau aggregates, and more recently in the formation of phase separated liquid droplets^11,35^, may also be important for the interaction of tau with microtubules, the most conspicuous tau interaction partner^16^. Here, by employing a combination of theory and experiments, we show that the interplay between tau–tau and tau–microtubule interactions drives the formation of multilayered tau condensates on the microtubule surface.

Over the past couple of decades, biomolecular condensation driven by phase separation has emerged as a fundamental paradigm for cellular organization^37–39^. A large body of experimental work has established that liquid–liquid phase separation serves as a pervasive organizing principle across molecular and cellular biology, underlying the formation of nucleoli and P-granules, as well as the assembly of diverse subcellular structures including heterochromatin^40,41^ and cytoskeletal filaments^42^ and has recently been extended to microtubule-associated proteins such as Kar9, which forms condensates at selected microtubule plus ends^43^. However, a comprehensive understanding of the biophysical principles governing biomolecular condensation inside cells remains elusive. A defining feature of the intracellular space is the prevalence of diverse surfaces, including lipid bilayers, DNA, and cytoskeletal networks, which serve as scaffolds for molecular interactions. The role of such surfaces in driving biomolecular condensation is still poorly understood and has only recently begun to attract significant attention. Notably, attractive surfaces can lower the nucleation barrier for condensation of proteins by promoting the formation of localized, multilayered liquid phases via a prewetting-like transition. For instance, recent studies have suggested prewetting phenomena to underlie proteins forming condensates on diverse surfaces such as DNA^37,38,44^ (see also Supplementary Fig. S8), lipid membranes^25,45^ and actin filaments^46^. One key criticism of the field of liquid-liquid phase separation in cells is that the concentrations required for phase separation *in vitro* are often substantially higher than physiologically relevant concentrations^47,48^. A prewetting transition provides a possible resolution to this discrepancy by allowing multilayered assemblies to form on intracellular surfaces even at physiologically relevant concentrations. For instance, the physiological concentration of tau in neurons has been estimated ∼0.5 – 2 µM^35,49^. Notably, tau binding to microtubules is already saturated at 0.5 µM in our in vitro assays, while in absence of crowding agents, in vitro tau condensation in solution, required more than 50 µM tau^35^. Thus, our work establishes microtubules as surfaces that can nucleate and organize condensates below saturation concentrations for phase separation in solution.

Tau condensation on microtubule surfaces, as observed in our experiments, happens below the threshold concentration for phase separation in solution. Unlike droplets in the solution, surface condensates are attached to their substrates and would dissolve upon removal of the surface. These assemblies are spatially constrained and limited in size to the regions where surface interactions favour their stability, as observed in our case. On flat surfaces, the thickness of a condensed layer grows indefinitely as the concentration approaches the saturation concentration for phase separation in solution from below^50^. In contrast, previous studies have shown that for surfaces with a sufficiently high curvature, such as a thin cylinder, the condensed layer thickness saturates to a finite value with increasing concentration. This phenomena is consistent with our observations, that the tau density on the microtubules saturate at higher concentrations. For a sufficiently thin cylinder, even above saturation concentrations, the surface condensation yields a wetting layer of finite thickness^30,31,51^. Such layers may eventually undergo a Plateau-Rayleigh instability, the same physical mechanism responsible for dew drop formation on spider silk, leading to a transition from uniform coatings of proteins on the microtubule surface into discrete droplets. This phenomenon was recently observed for TPX2, a protein that reorganizes from a uniform coating into regularly spaced condensates that locally concentrate branching factors, including augmin and γ-TuRC, thereby creating discrete reaction hubs from which branched microtubules nucleate^52,53^. A similar mechanism has recently been proposed for the microtubule plus-end tracking protein EB3, where surface condensation generates a wetting layer at growing microtubule ends and a subsequent Plateau–Rayleigh instability limits the length of the resulting EB3 comet^54^. Another recent study showed the role of capillary forces, generated by MAP condensates, in microtubule bundling^53^.

The physical nature of the microtubule-tau interaction has thus far remained unclear. Previous observations of dense tau patches on microtubules have been referred to as condensates^17^. A recent review further suggested, based on stoichiometry of tau:tubulin, that the observed tau patches could be surface-bound condensates, and are likely multilayered in nature^55^. Here, we develop theory describing the formation of condensates through a prewetting-like mechanism and provide experimental evidence of their multilayered nature. While we cannot completely rule out that high-turnover tau molecules, comprising the outer layers of the condensates, are in direct weak contact with the microtubule lattice, e.g. via one of their microtubule-binding repeats, a systematic comparison between our theory and experiments suggests that tau–tau interactions play an important role in governing how tau molecules bind to microtubules. Our picture is further supported by the observation that in presence of crowding agents tau can undergo phase separation in solution, at concentrations of around 5-10 μM^11,26,35^, through tau–tau interactions. Tau droplets have also been shown to recruit tubulin monomers and promote microtubule filament formation^11^, consistent with our observation that tau condensates assembled on microtubule surfaces can similarly incorporate tubulin.

Importantly, experiments presented in this and previous studies suggest that the two distinct populations of tau (*please refer to the graphical abstract, below*) within tau condensates have different functional properties. Numerous previous experiments pointed out that the cooperatively adsorbed, low turnover tau, has selectively restrictive function: it prevents the binding of specific microtubule-associated proteins, such as kinesin-1 or katanin, to the microtubule surface, while allowing binding of other proteins, such as kinesin-3 or dynein^6,16–19^. Our current study shows that the high turnover tau layer, by contrast, has the ability to selectively recruit and thus locally enrich tau interactors, such as soluble tubulin, microtubules, actin and RNA to the microtubule surface. Such local enrichment can have functional consequences, as exemplified here by enhanced microtubule growth by recruited soluble tubulin. Effectively, such tau condensate can be seen as a membrane-less compartment establishing a specific microenvironment. These observations are in line with recent data showing that tau and tubulin can form a viscoelastic intervening network, which can cross-bridge microtubules^56^, that tau enhances microtubule lattice repair by increasing the incorporation of tubulin into the shaft of pre-existing microtubules^57^, and that tau-DNA condensates can interact with microtubules^24^. Furthermore, while tau is often referred to as a microtubule-stabilizing or microtubule-bundling protein, multiple reports challenge this prevailing view ^58–61^. Our results indicate that a prewetting transition, dictated by tau-tau and tau-microtubule interactions might act as a switch for these processes, potentially offering reconciling view to these observations. While the cooperative formation of the adsorbed layer depends on the conformation of tubulin and occurs preferentially on compacted microtubule lattice, the high-turnover condensate can form on both extended and compacted lattices. Recent observation of extended tubulin lattices in axons^62^ thus suggests that tau in the axon might bind to microtubules predominantly with high turnover, in accordance with previous in vivo observations^58^, potentially forming a specific microenvironment around the axonal microtubule bundles.

Overall, our findings suggest a key role of the microtubule surface in the formation of tau condensates: a pre-wetting like transition lowers the tau concentration required for tau phase separation by two orders of magnitude (comparing the 0.5 µM tau used in the present study with the >50 µM tau critical concentration required for in vitro reconstitution of liquid droplet formation in the absence of crowding agents^35^, well into the physiological range estimated in neurons (∼0.5 −2 µM). We demonstrate functional consequences of the enwrapping of filament surfaces in liquid films, namely the crosslinking of filaments, as well as the local recruitment of RNA and, selectively, of client proteins, to the close microtubule vicinity. We anticipate our description and demonstration of pre-wetting like transitions to provide a theoretical framework applicable to various biological surfaces beyond the cytoskeleton.

## Data availability

Source data are provided with this paper.

## Acknowledgements

We thank K. Konečná, C. de Brito and V. Rumlová for technical support. This project has received funding from the European Research Council under the European Union’s Horizon Europe research and innovation program (grant no. 101071583 (TubulinCode)); European Molecular Biology Organization postdoctoral fellowship (ALTF 1069-2024 to V.S.); the Czech Science Foundation (25-16671S to M. Braun) and S.C. is supported by the Ramalingaswami Re-entry Fellowship (BT/HRD/35/02/2006), a re-entry scheme of the Department of Biotechnology, Ministry of Science and Technology, Government of India. The authors acknowledge Imaging Methods Core Facility at BIOCEV – a member of Euro-BioImaging ERIC and Czech-BioImaging national infrastructure supported by MEYS LM2023050 grant – for their support & assistance in TIRF microscopy and Charles University – Viničná Microscopy Core Facility (RRID:SCR_026602), Faculty of Science, Czech-BioImaging large RI project LM2023050, e-INFRA CZ project (ID:90254). We acknowledge CF ProtProd of CIISB, Instruct-CZ Centre, supported by MEYS CR (LM2023042) and European Regional Development Fund-Project „Innovation of Czech Infrastructure for Integrative Structural Biology” (No. CZ.02.01.01/00/23_015/0008175).

## Author contributions

S.C., M.B. and Z.L. conceptualized the manuscript. E.L., T.H., V.S., J.K. and M.D.Z. developed the experimental methods. A.N. and S.C. developed the theory. A.N. and S.C. performed simulations. E.L., T.H., V.S., J.K., M.D.Z and A.B. performed in vitro TIRF experiments. S.K. purified HeLa tubulin. E.L., T.H., J.K. and M.D.Z. analyzed the experimental data and A.N. analyzed the theoretical data. S.K. and C.J. provided resources. E.L., A.N., Z.L., M.B. and S.C. wrote the manuscript, which was reviewed and edited by M.D.Z. and V.S. E.L., A.N., V.S. and M.B. were responsible for visualization. L.L., Z.L., M.B. and S.C. supervised the project. V.S., Z.L, M.B. and S.C. acquired funding.

## Ethics declarations

The authors declare no competing interests.

## Methods

### 1. Protein constructs and purification

For the in vitro experiments, tau N-terminally-tagged with mNeonGreen (tau-mNG, 2N4R tau; NM_005910.6; subcloned into an expression vector based on pET11Kan-N-HIS6-3C-mNeon-Green) was expressed in E. coli BL21(DE3)-RIPL cells. Cells were grown at 30 °C to an OD600 of 0.5–0.6, after which protein expression was induced with 0.1 mM IPTG and cultures were incubated overnight at 16 °C. Bacterial pellets were resuspended in 50 mL lysis buffer (50 mM HEPES pH 7.4, 300 mM KCl, 2 mM MgCl_2_, 0.1% Tween 20, 5% glycerol, 1 mM DTT, 0.1 mM ATP, 10 mM β-mercaptoethanol, 1× protease inhibitor cocktail and Benzonase, 25 U), sonicated on ice (10 min, 2 s ON/4 s OFF) and centrifuged at 180,000 × g for 60 min at 4 °C. The soluble fraction was incubated with HisTrap Ni-NTA agarose resin (XF340049, Thermo Scientific) for 2 h at 4 °C with slow rotation. Beads were washed with 20 mL Wash Buffer I (50 mM HEPES pH 7.4, 300 mM KCl, 0.1% Tween 20, 5% glycerol, 1 mM DTT, 0.1 mM ATP and 30 mM imidazole), followed by 10 mL Wash Buffer II (Wash Buffer I supplemented with 60 mM imidazole). Proteins were eluted using Elution Buffer (Wash Buffer I supplemented with 250 mM imidazole). Eluted fractions were further purified using Strep-Tactin XT affinity purification (wash buffer: 50 mM HEPES pH 7.4, 300 mM KCl, 2 mM MgCl_2_, 1 mM EGTA, 0.1% 20, 5% glycerol, 1 mM DTT and 0.1 mM ATP) and eluted with wash buffer supplemented with 50 mM biotin. Eluted proteins were dialyzed overnight against dialysis buffer (50 mM HEPES pH 7.4, 300 mM KCl, 0.1% Tween 20, 5% glycerol and 30 mM imidazole) in 12 kDa cut-off dialysis tubing in the presence of 3C protease to cleave the 6×His and Strep tags. Following dialysis, the sample was incubated with Ni-NTA agarose resin to remove His-tagged 3C protease, and the flow-through containing purified protein was collected. Purified proteins were concentrated using VivaSpin-10 kDa-HYProtein, and protein concentration was determined by NanoDrop absorbance measurements at 280 nm. Purified proteins were flash-frozen in liquid nitrogen and stored at −80 °C. All purification procedures were performed at 4 °C. Tau N-terminally tagged with meGFP (tau-meGFP) was expressed in insect cells and purified as described previously^11^.

### 2. HeLa cell tubulin isolation

#### HeLa cells culture and lysate preparation

HeLa S3 cells were expanded for 7 days in spinner bottles containing 1 L of DMEM supplemented with 10% heat-inactivated foetal bovine serum, 2 mM L-glutamine, and 1× penicillin–streptomycin, and maintained under stirring in a humidified cell-culture incubator at 37 °C and 5% CO₂. A detailed protocol for adapting and amplifying cells for suspension culture is provided in Souphron et al., 2019^63^. Suspension cultures were harvested and centrifuged at 250×g for 15 min at room temperature. The supernatant was discarded carefully, and the pellet was washed in 30 ml of PBS and centrifuged at 250×g for 8 min at RT. After discarding the supernatant, the cell pellet was resuspended in an equal volume of ice-cold lysis buffer (BRB80, pH 6.8, supplemented with 1 mM β-mercaptoethanol, 1 mM PMSF, and 1× home-made protease inhibitor cocktail containing 20 μg/ml leupeptin, 20 μg/ml aprotinin and 20 μg/ml 4-(2-aminoethyl)-benzene sulfonyl fluoride). Cells were lysed using a French press, and the lysate was clarified by ultracentrifugation at 112,000×g for 30 min at 4 °C to obtain the first supernatant.

#### Tubulin purification from HeLa cells in suspension culture

The tubulin purification was performed using successive cycles of polymerisation and depolymerisation, as detailed in Souphron et al., 2019^63^. In brief, the lysate supernatant was collected with a syringe fitted with a long needle and was immediately incubated on ice. The volume of the supernatant was recorded. For the first polymerisation-depolymerisation cycle, the supernatant was supplemented with 1 mM GTP and 30% (v/v) prewarmed glycerol and incubated for 30 min in a water bath at 30 °C. Following incubation, the sample was centrifuged in a prewarmed rotor at 112,000 × g for 30 min at 30 °C to pellet polymerised microtubules and associated proteins. The pellet was depolymerised on ice for 5 min in ice-cold BRB80 corresponding to 1/60 of the initial recorded supernatant volume, followed by pipetting up and down for 25 to 35 min, also on ice. The resuspended pellet was then centrifuged in a pre-cooled rotor at 112,000 × g for 20 min at 4 °C. A second polymerisation was then performed by mixing the obtained supernatant 1:1 (v/v) with 1 M PIPES, pH 6.8, yielding 0.5 M PIPES final concentration, followed by the addition of 1 mM GTP and glycerol to 30% (v/v) and incubating for 30 min at 30 °C. This was followed by a depolymerisation step in which 1/100 of the supernatant volume of ice-cold BRB80 was added. The microtubule pellet was depolymerised on ice and centrifuged at 112,000 × g for 20 min at 4 °C. The third polymerisation was performed similarly to the first cycle. The final microtubule pellet was depolymerised on ice for 15 min with ice-cold BRB80 at 1/300 of the initial clarified lysate volume and centrifuged at 112,000 × g for 20 min at 4 °C to yield soluble tubulin in the supernatant. Following this step, tubulin concentration was estimated by measuring absorbance at A280 with a NanoDrop (ND-1000 spectrophotometer; Thermo Scientific; MW = 110 kDa; ε = 115,000 M⁻¹/cm⁻¹), and purity was assessed by Coomassie-stained SDS–PAGE. Purified tubulin was aliquoted and snap-frozen in liquid nitrogen and stored at −80 °C until use.

### 3. Microtubule Assembly

Porcine brain tubulin was isolated using the high-molarity PIPES procedure as previously described^64^. Biotin-labeled and HiLyte647-labeled tubulin were purchased from Cytoskeleton Inc. (T333P and TLL488M). HeLa tubulin was isolated from HeLa S3 cells as described above.

*Taxol-stabilized microtubules* were polymerized from 4 mg/mL tubulin in BRB80 buffer (80 mM PIPES, 1 mM EGTA and 1 mM MgCl_2_, pH 6.9) supplemented with 4 mM MgCl_2_, 1 mM GTP (Jena Bioscience, NU-1012) and 5% DMSO for 30 min at 37 °C. Polymerized microtubules were diluted in 100 µL BRB80T (BRB80 supplemented with 10 µM taxol) and pelleted by centrifugation for 30 min at room temperature using either a Hettich Universal 320R centrifuge (21,380 × g, rotor 1420-A) or a Beckman Microfuge (18,000 × g). Pellets were resuspended in BRB80T, stored at room temperature and used within one week of preparation.

*GMPCPP-stabilized microtubules* were polymerized from 4 mg/mL tubulin (1:50 biotin-labeled to unlabeled tubulin) for 4 h at 37 °C in BRB80 supplemented with 1 mM MgCl_2_ and 1 mM GMPCPP (Jena Bioscience, NU-405). Polymerized microtubules were centrifuged for 30 min at room temperature using either a Hettich Universal 320R centrifuge (21,380 × g, rotor 1420-A) or a Beckman Microfuge (18,000 × g). Pellets were resuspended in BRB80, stored at room temperature and used within one week of preparation.

*GDP microtubules (stabilized with GMPCPP caps)* were polymerized from 4 mg/mL tubulin (1:50 biotin-labeled to unlabeled tubulin) in BRB80 supplemented with 4 mM MgCl_2_, 1 mM GTP (Jena Bioscience, NU-1012) and 5% DMSO for 90 min at 37 °C. Polymerized microtubules were centrifuged for 30 min at 21,380 × g at room temperature. Pellets were carefully resuspended in 35 µL of prewarmed BRB80 supplemented with 1.25 mM GMPCPP and 1.25 mM MgCl_2_, followed by addition of 0.7 µL HiLyte647-labeled tubulin (4 mg/mL). Samples were incubated for 10 min at 37 °C and subsequently kept at room temperature. GDP-microtubules were used on the same day they were prepared.

### 4. F-actin assembly and RNA preparation

Lyophilized rabbit muscle actin (Cytoskeleton, AKL99) was resuspended and stored according to the manufacturer’s instructions. Actin filaments (final concentration 400 mg/mL) were polymerized in actin polymerization buffer (5 mM Tris-HCl pH 7.8, 0.2 mM CaCl_2_, 50 mM KCl, 2 mM MgCl_2_ and 1 mM ATP) supplemented with rhodamine-phalloidin (10 mM final concentration) for filament stabilization and labeling. Polymerization was carried out overnight at 4 °C, and stabilized filaments were used within two weeks. Prior to experiments, actin filaments were diluted and mechanically fragmented as described below. RNA oligonucleotides (poly-A-RNA-Cy5; SC1518) were synthesized by GenScript and diluted as indicated below.

### 5. Total internal reflection fluorescence microscopy (TIRF)

Total internal reflection fluorescence (TIRF) microscopy was performed using either a Nikon Ti-E inverted microscope equipped with an H-TIRF module or a Zeiss Elyra PS.1 microscope. Imaging on the Nikon Ti-E was performed using a 60× oil-immersion objective (NA 1.49, Apo TIRF; Nikon) and sCMOS cameras (Hamamatsu Photonics). Microtubules were visualized by interference reflection microscopy (IRM), whereas fluorescently labeled proteins were imaged using EGFP, mCherry and Cy5 filter sets or a quad-band filter set (405, 488, 561 and 640 nm). The microscope was controlled using NIS-Elements AR software (v. 5.20). Imaging on the Zeiss Elyra PS.1 microscope was performed using a 100×/1.46 oil-immersion objective and an EMCCD Andor PALM camera. Fluorescently labeled microtubules and tau-mNG were visualized using 647 nm and 488 nm lasers, respectively. The microscope was controlled using ZEN Black software. All experiments were performed at room temperature.

TIRF flow chambers were assembled using thin parafilm strips and HMDS-treated glass coverslips (22 × 22 mm and 18 × 18 mm; Corning). Chambers were incubated with anti-biotin antibodies (20 µg/mL; Sigma Aldrich, B3640) or anti-β-tubulin antibodies (20 µg/mL; Sigma, T7816) in PBS for 5 min and subsequently blocked with 1% Pluronic F127 (Sigma Aldrich, P2443) for at least 30 min. Chambers were washed with 40 µL BRB80T prior to experiments.

### 6. TIRF assays

#### In vitro tau–microtubule binding

mNeonGreen–tau was diluted in TIRF assay buffer (50 mM HEPES pH 7.4, 1 mM EGTA, 10 µM taxol, 2 mM MgCl_2_, 75 mM KCl, 10 mM DTT, 0.5 mg/mL casein, 1 mM Mg-ATP, 20 mM D-glucose and 0.1% Tween 20). Following immobilization of microtubules on the coverslip surface, tau was diluted to the indicated final concentration in TIRF imaging buffer (assay buffer supplemented with 0.22 mg/mL glucose oxidase and 20 µg/mL catalase) and introduced into the imaging chamber at twice the chamber volume. Tau binding was monitored by time-lapse imaging.

#### Multiple tau addition and removal

mNeonGreen–tau was diluted in TIRF imaging buffer to a final concentration of 5 nM and incubated on surface-immobilized taxol-stabilized microtubules for 10 min. Tau was removed by washing the chamber three times with 20 µL TIRF assay buffer, followed by a wash with assay buffer containing elevated ionic strength (125 mM KCl), and two further washes with TIRF assay buffer. Following tau removal, 5 nM tau in TIRF imaging buffer was reintroduced into the chamber. This procedure was repeated three times and tau binding was monitored by time-lapse imaging.

#### Tau binding to porcine brain and HeLa microtubules

Taxol-stabilized microtubules polymerized from porcine brain tubulin (1:5 HiLyte647-labeled to unlabeled tubulin) or HeLa tubulin (unlabeled) were prepared as described above. Microtubules were mechanically sheared using a Hamilton syringe to generate short seeds, mixed at a 1:1 ratio and incubated overnight at room temperature to allow formation of long hybrid microtubules. Microtubules were immobilized on the coverslip surface, and diluted tau was introduced into the chamber. Tau binding was monitored by time-lapse imaging, and the fraction of microtubule length covered by tau condensates was quantified.

#### Tau unbinding in time

Taxol-stabilized, GDP- or GMPCPP-stabilized microtubules were introduced into the chamber and incubated with tau diluted in TIRF imaging buffer at concentrations indicated in the corresponding figure legends. Samples were allowed to equilibrate for 3 min prior to tau removal by washing with 30 µL TIRF imaging buffer. Tau dissociation was monitored by time-lapse imaging. Experiments with taxol-stabilized microtubules were performed in the presence of 10 µM taxol, whereas GDP- and GMPCPP-stabilized microtubules were performed in taxol-free buffer.

#### Recruitment of tubulin and short microtubules to microtubule-associated tau condensates

Taxol-stabilized, GDP-lattice or GMPCPP-lattice microtubules were introduced into imaging chambers, followed by addition of tau diluted in BRB80-TIRF imaging buffer (BRB80 supplemented with 10 mM DTT, 10 µM taxol, 0.05 mg/mL casein, 1 mM Mg-ATP, 20 mM D-glucose, 0.1% Tween 20, 0.22 mg/mL glucose oxidase and 20 µg/mL catalase) to final concentrations of 5 nM or 500 nM. Tau-free conditions were included as controls. Following a 3 min equilibration period, binding partners were added. HiLyte647-labeled tubulin was added at final concentrations of 90, 180 or 900 nM. For recruitment of short microtubules, taxol-stabilized microtubules were polymerized from a 5:1 mixture of unlabeled and HiLyte647-labeled tubulin, mechanically sheared using a Hamilton syringe to generate short seeds and introduced into the imaging chamber.

#### Tau interaction with dynamic microtubules\

GMPCPP-stabilized microtubule seeds were introduced into imaging chambers and unbound seeds were removed by washing with BRB80. Chambers were equilibrated with BRB80-TIRF assay buffer (BRB80 supplemented with 10 mM DTT, 0.05 mg/mL casein, 1 mM Mg-ATP, 20 mM D-glucose and 0.1% Tween 20). For each tau condition (0, 5 and 500 nM), three tubulin concentrations were tested (3.6, 7.2 and 10.8 µM). Tubulin and tau were diluted in polymerization buffer (BRB80 assay buffer supplemented with 0.1% methylcellulose, 2.5 mM GTP, 10 mM MgCl_2_, 0.22 mg/mL glucose oxidase and 20 µg/mL catalase), introduced into the chamber and microtubule growth dynamics were recorded.

#### RNA and F-actin recruitment assays

Pre-polymerized taxol-stabilized microtubules were introduced into imaging chambers and washed with BRB80-TIRF assay buffer to remove unbound microtubules. Tau-mNG was diluted in BRB80-TIRF imaging buffer to final concentrations of 10 nM or 1 µM, or omitted in control conditions. Samples were incubated for at least 5 min to allow tau binding to reach equilibrium. For F-actin recruitment experiments, prepolymerized, phalloidin-stabilized, rhodamine-labeled actin filaments (final actin concentration 4 µg/mL) were mixed with tau-mNG at the indicated concentrations and immediately introduced into the chamber for TIRF imaging. For RNA recruitment experiments, RNA oligonucleotides were diluted to a final concentration of 5 nM, mixed with tau-mNG at the indicated concentrations and incubated on ice for 10 min prior to introduction into the imaging chamber. Afterwards, the mix was introduced to the chamber for TIRF imaging at room temperature.

### 7. TIRF image analysis

Microscopy data were analyzed using ImageJ/Fiji (version 2.16.0/1.54q). Kymographs of individual microtubules were generated using the KymographBuilder plugin by drawing segmented lines along the microtubule lattice. Data analysis and graph generation were performed using Julia (version 1.10.9) or GraphPad Prism (Prism 11.0.2).

#### Tau, tubulin, actin and RNA density analysis

Protein and RNA fluorescence intensity on microtubules was quantified by drawing a line along the entire microtubule lattice and measuring the mean grey value. Background fluorescence was measured using the same line placed in an adjacent microtubule-free region and subtracted from the corresponding microtubule-associated signal.

#### The fraction of microtubule length covered by tau condensates

The fraction of microtubule length covered by tau condensates was determined by measuring the length of tau condensates on individual microtubules and dividing the summed condensate length by the total microtubule length within the respective field of view.

#### Estimation of the tau-unbinding time

To estimate unbinding times, we quantified the decay of tau density in a defined region on microtubule following buffer exchange that removed tau from solution. When necessary, time series were drift-corrected with the FIJI plugin ‘image stabilizer’. Individual microtubules were traced using segmented lines, and the mean fluorescence intensity of tau was measured along each filament at every frame (500 ms intervals) after tau removal from solution. Background fluorescence, obtained from adjacent microtubule-free regions, was subtracted from the corresponding microtubule-associated signal to yield background-corrected tau intensity traces.

#### Dynamic microtubules analysis

For microtubule dynamics experiments, growth rates (nm/s) and catastrophe frequencies (s⁻¹) were determined from kymographs. Growth rates were calculated from the slopes of growth events in the kymographs and expressed in nm/s. Catastrophe frequency was calculated as the number of catastrophe events divided by the total time spent in the growth phase. Only microtubules that remained visible throughout the entire observation period were included in the analysis.

## References

1. Janke, C. & Magiera, M. M. The tubulin code and its role in controlling microtubule properties and functions. Nat. Rev. Mol. Cell Biol. 21, 307–326 (2020).

2. Alushin, G. M. et al. High-resolution microtubule structures reveal the structural transitions in αβ-Tubulin upon GTP hydrolysis. Cell 157, 1117–1129 (2014).

3. de Jager, L. et al. StableMARK-decorated microtubules in cells have expanded lattices. J. Cell Biol. 224, e202206143 (2024).

4. Shima, T. et al. Kinesin-binding-triggered conformation switching of microtubules contributes to polarized transport. J. Cell Biol. 217, 4164–4183 (2018).

5. Peet, D. R., Burroughs, N. J. & Cross, R. A. Kinesin expands and stabilizes the GDP-microtubule lattice. Nat. Nanotechnol. 13, 386–391 (2018).

6. Siahaan, V. et al. Microtubule lattice spacing governs cohesive envelope formation of tau family proteins. Nat. Chem. Biol. 18, 1224–1235 (2022).

7. Liu, H. & Shima, T. Preference of CAMSAP3 for expanded microtubule lattice contributes to stabilization of the minus end. Life Sci. Alliance 6, e202201714 (2023).

8. Paquette, A. L. et al. Competition for microtubule lattice spacing between a microtubule expander and compactor. Curr. Biol. 35, 4442–4452.e4 (2025).

9. Kellogg, E. H. et al. Insights into the Distinct Mechanisms of Action of Taxane and Non-Taxane Microtubule Stabilizers from Cryo-EM Structures. J. Mol. Biol. 429, 633–646 (2017).

10. Bodakuntla, S., Jijumon, A. S., Villablanca, C., Gonzalez-Billault, C. & Janke, C. Microtubule-Associated Proteins: Structuring the Cytoskeleton. Trends Cell Biol. 29, 804–819 (2019).

11. Hernández-Vega, A. et al. Local Nucleation of Microtubule Bundles through Tubulin Concentration into a Condensed Tau Phase. Cell Rep. 20, 2304–2312 (2017).

12. Rai, S. K., Savastano, A., Singh, P., Mukhopadhyay, S. & Zweckstetter, M. Liquid–liquid phase separation of tau: From molecular biophysics to physiology and disease. Protein Sci. 30, 1294–1314 (2021).

13. Boyko, S. & Surewicz, W. K. Tau liquid–liquid phase separation in neurodegenerative diseases. Trends Cell Biol. 32, 611–623 (2022).

14. Islam, M. et al. Tau liquid-liquid phase separation: At the crossroads of tau physiology and tauopathy. J. Cell. Physiol. 239, e30853 (2024).

15. Hinrichs, M. H. et al. Tau Protein Diffuses along the Microtubule Lattice*. J. Biol. Chem. 287, 38559–38568 (2012).

16. Siahaan, V. et al. Kinetically distinct phases of tau on microtubules regulate kinesin motors and severing enzymes. Nat. Cell Biol. 21, 1086–1092 (2019).

17. Tan, R. et al. Microtubules gate tau condensation to spatially regulate microtubule functions. Nat. Cell Biol. 21, 1078–1085 (2019).

18. Monroy, B. Y. et al. Competition between microtubule-associated proteins directs motor transport. Nat. Commun. 9, 1487 (2018).

19. Zhernov, I., Diez, S., Braun, M. & Lansky, Z. Intrinsically Disordered Domain of Kinesin-3 Kif14 Enables Unique Functional Diversity. Curr. Biol. 30, 3342–3351.e5 (2020).

20. Bonn, D. & Ross, D. Wetting transitions. Rep. Prog. Phys. 64, 1085 (2001).

21. Zhao, X., Bartolucci, G., Honigmann, A., Jülicher, F. & Weber, C. A. Thermodynamics of wetting, prewetting and surface phase transitions with surface binding. New J. Phys. 23, 123003 (2021).

22. Cahn, J. W. Critical point wetting. J. Chem. Phys. 66, 3667–3672 (2008).

23. Chen, X., Jia, B., Zhu, S. & Zhang, M. Phase separation-mediated actin bundling by the postsynaptic density condensates. eLife 12, e84446 (2023).

24. Park, C. et al. Tau condensation on DNA mediates microtubule attachment suggesting a mitotic role for centromere-localized tau. Nat. Commun. 17, 1128 (2026).

25. Shaikh, J. et al. Membrane interfacial potential governs surface condensation and fibrillation of α-Synuclein in neurons. Nat. Commun. 17, 4247 (2026).

26. Boyko, S., Surewicz, K. & Surewicz, W. K. Regulatory mechanisms of tau protein fibrillation under the conditions of liquid–liquid phase separation. Proc. Natl. Acad. Sci. 117, 31882–31890 (2020).

27. Genova, M. et al. Tubulin polyglutamylation differentially regulates microtubule-interacting proteins. EMBO J. 42, EMBJ2022112101 (2023).

28. Boucher, D., Larcher, J.-C., Gros, F. & Denoulet, P. Polyglutamylation of Tubulin as a Progressive Regulator of in Vitro Interactions between the Microtubule-Associated Protein Tau and Tubulin. Biochemistry 33, 12471–12477 (1994).

29. Liu, H., Yamaguchi, H., Kikkawa, M. & Shima, T. Heterogeneous local structures of the microtubule lattice revealed by cryo-ET and non-averaging analysis. Preprint at 10.1101/2024.04.30.591984 (2024).

30. Gelfand, M. P. & Lipowsky, R. Wetting on cylinders and spheres. Phys. Rev. B 36, 8725–8735 (1987).

31. Bieker, T. & Dietrich, S. Wetting of curved surfaces. Phys. Stat. Mech. Its Appl. 252, 85–137 (1998).

32. Kanaan, N. M. & Grabinski, T. Neuronal and Glial Distribution of Tau Protein in the Adult Rat and Monkey. Front. Mol. Neurosci. 14, (2021).

33. Cabrales Fontela, Y., et al. Multivalent cross-linking of actin filaments and microtubules through the microtubule-associated protein Tau. Nat. Commun. 8, 1981 (2017).

34. He, H. J. et al. The proline-rich domain of tau plays a role in interactions with actin. BMC Cell Biol. 10, 81 (2009).

35. Wegmann, S. et al. Tau protein liquid–liquid phase separation can initiate tau aggregation. EMBO J. 37, EMBJ201798049 (2018).

36. Hochmair, J. et al. Molecular crowding and RNA synergize to promote phase separation, microtubule interaction, and seeding of Tau condensates. EMBO J. 41, EMBJ2021108882 (2022).

37. Hyman, A. A., Weber, C. A. & Jülicher, F. Liquid-Liquid Phase Separation in Biology. Annu. Rev. Cell Dev. Biol. 30, 39–58 (2014).

38. Banani, S. F., Lee, H. O., Hyman, A. A. & Rosen, M. K. Biomolecular condensates: organizers of cellular biochemistry. Nat. Rev. Mol. Cell Biol. 18, 285–298 (2017).

39. Alberti, S. et al. Current practices in the study of biomolecular condensates: a community comment. Nat. Commun. 16, 7730 (2025).

40. Larson, A. G. et al. Liquid droplet formation by HP1α suggests a role for phase separation in heterochromatin. Nature 547, 236–240 (2017).

41. Strom, A. R. et al. Phase separation drives heterochromatin domain formation. Nature 547, 241–245 (2017).

42. Wiegand, T. & Hyman, A. A. Drops and fibers — how biomolecular condensates and cytoskeletal filaments influence each other. Emerg. Top. Life Sci. 4, 247–261 (2020).

43. Meier, S. M. et al. Multivalency ensures persistence of a +TIP body at specialized microtubule ends. Nat. Cell Biol. 25, 56–67 (2023).

44. Morin, J. A. et al. Sequence-dependent surface condensation of a pioneer transcription factor on DNA. Nat. Phys. 18, 271–276 (2022).

45. Rouches, M., Veatch, S. L. & Machta, B. B. Surface densities prewet a near-critical membrane. Proc. Natl. Acad. Sci. 118, e2103401118 (2021).

46. Wiegand, T., et al. Actin polymerization counteracts prewetting of N-WASP on supported lipid bilayers. Proc. Natl. Acad. Sci. 121, e2407497121 (2024).

47. Musacchio, A. On the role of phase separation in the biogenesis of membraneless compartments. EMBO J. 41, e109952 (2022).

48. Volkov, V. A. & Akhmanova, A. Phase separation on microtubules: from droplet formation to cellular function? Trends Cell Biol. 34, 18–30 (2024).

49. Drubin, D. G., Feinstein, S. C., Shooter, E. M. & Kirschner, M. W. Nerve growth factor-induced neurite outgrowth in PC12 cells involves the coordinate induction of microtubule assembly and assembly-promoting factors. J. Cell Biol. 101, 1799–1807 (1985).

50. Cahn, J. W. Critical point wetting. J. Chem. Phys. 66, 3667–3672 (1977).

51. Levinson, P., Jouffroy, J. & Brochard, F. Wetting transition for a thin cylinder. J. Phys. Lett. 46, 21–26 (1985).

52. Setru, S. U. et al. A hydrodynamic instability drives protein droplet formation on microtubules to nucleate branches. Nat. Phys. 17, 493–498 (2021).

53. Gouveia, B. et al. Capillary bundling of microtubules by condensates. bioRxiv 2026.06.19.733462 (2026) doi:10.64898/2026.06.19.733462.

54. Schaer, J. et al. Phase Separation and a Hydrodynamic Instability Localize Proteins at Growing Microtubule Ends.

55. Mitchison, T. J. Beyond Langmuir: surface-bound macromolecule condensates. Mol. Biol. Cell 31, 2502–2508 (2020).

56. Kohl, P. A. et al. Complexes of tubulin oligomers and tau form a viscoelastic intervening network cross-bridging microtubules into bundles. Nat. Commun. 15, 2362 (2024).

57. Biswas, S. et al. Tau accelerates tubulin exchange in the microtubule lattice. Nat. Phys. 21, 1616–1628 (2025).

58. Bakota, L. & Brandt, R. Why kiss-and-hop explains that tau does not stabilize microtubules and does not interfere with axonal transport (at physiological conditions). Cytoskeleton 81, 47–52 (2024).

59. Baas, P. W. & Qiang, L. Tau: It’s Not What You Think. Trends Cell Biol. 29, 452–461 (2019).

60. Qiang, L. et al. Tau Does Not Stabilize Axonal Microtubules but Rather Enables Them to Have Long Labile Domains. Curr. Biol. CB 28, 2181–2189.e4 (2018).

61. Janning, D. et al. Single-molecule tracking of tau reveals fast kiss-and-hop interaction with microtubules in living neurons. Mol. Biol. Cell 25, 3541–3551 (2014).

62. Zehr, E. A., Sun, S., Sarbanes, S. L. & Roll-Mecak, A. Microtubules in the axon are GDP bound but adopt a stable GTP-like expanded state. Nat. Struct. Mol. Biol. 33, 631–640 (2026).

63. Souphron, J. et al. Purification of tubulin with controlled post-translational modifications by polymerization–depolymerization cycles. Nat. Protoc. 14, 1634–1660 (2019).

64. Castoldi, M. & Popov, A. V. Purification of brain tubulin through two cycles of polymerization–depolymerization in a high-molarity buffer. Protein Expr. Purif. 32, 83–88 (2003).

