## Supplementary material for "Surface-induced tau condensation generates a selective microenvironment around microtubules": Supplemetary information and figures

**Supplementary Information**

### 1. Lattice-gas model for tau condensation on microtubules

#### 1.1. Model overview

In order to study the interaction of tau molecules with microtubules, we use a minimal lattice gas model. In our model, tau molecules are present at the vertices of a three-dimensional cubic lattice with $L$x$M$x$L$ sites and periodic boundary conditions. The occupancy at site $i$ is given by $n_i = 0$ or 1, depending on whether it is unoccupied or a tau molecule is present, respectively. The microtubule is represented as a cylindrical surface of length $M$ lattice units, with 12 binding sites radially around it at each position (Supplementary Fig.S1A). This is motivated by, and approximated to, the fact that microtubules have 13 protofilaments on average[2]. Tau molecules interact favourably with each other, forming liquid droplets in the solution[11], and also interact with microtubules directly[65,66]. We incorporate these interactions in our framework via the following Hamiltonian:

$$H = -J_b \sum_{\langle i,j\rangle \in \text{bulk}} n_i\, n_j - J_s \sum_{\langle i,j\rangle \in \text{surf}} n_i\, n_j - \sum_{i \in \text{surf}} h_i\, n_i - \mu \sum n_i \quad (1)$$

The first term accounts for tau-tau interaction in solution, characterized by the coupling parameter $J_b$ (referred to as bulk coupling parameter). The sum is over all nearest neighbour pairs in the bulk, where each tau molecule can interact with other molecules at its 26 nearest neighbouring sites (6 along edges, 12 along face diagonals and 8 along body diagonals). In our model, tau can interact with the microtubule in two ways: first, individual molecules can bind to the surface at position $i$ with the binding energy $h_i$, as shown in the third term in the Hamiltonian. Second, tau molecules bound adjacently on the microtubule interact through the surface coupling parameter $J_s$. This is represented in the second term of the Hamiltonian and accounts for the binding cooperativity of tau. The final term, parametrized by the bulk chemical potential $\mu$, allows the exchange of molecules with an effective reservoir while keeping the average concentration of tau in the solution fixed. Similar lattice-based models have previously been used to study phase separation, as well as to study wetting phenomena[67–69].

#### 1.2. Simulation scheme

To evolve the system to equilibrium, we simulate it using the kinetic Monte Carlo method, where each step is performed as follows: we randomly choose between particle displacement, particle insertion into an empty site, and particle deletion from an occupied site, with probabilities of 0.5, 0.25, and 0.25, respectively. In a displacement step, a randomly chosen particle can be moved to any one of its neighbouring empty sites. For particle insertion, we choose a site at random and add a particle if it is empty. Particle deletion is carried out in a similar manner. All particle insertion and deletion moves are done only within a padded region 3 sites from the boundaries parallel to the microtubule, sufficiently far from the surface. This ensures that the local densities are preserved near the surface and that the proper kinetics of condensate growth are obtained. All moves are accepted with a probability given by the Metropolis condition, $p_{acc} = min\ [1, exp(-\beta \Delta H)]$, where $\Delta H$ is the change in the energy obtained from Eq.1. The system is initialized randomly with an initial density $\phi$=0.05 in the bulk. We simulate the system for $10^4$ to $10^5$ Monte Carlo sweeps for equilibration, with each sweep consisting of $L$x$M$x$L$ moves. For cases where we require only the equilibrium states, we relax the nearest-neighbour constraint on the displacement step and allow the particle to be globally displaced anywhere within the system.

##### 1.3. Identification of the phase coexistence region in the solution

The concentration of tau in all the experiments is below the phase separation threshold (undersaturated regime). We first carry out simulations without the microtubule present to identify the bulk coexistence region. For this, we work in the canonical (NVT) ensemble, where the bulk volume fraction $\phi$ is specified. The last term in Eq.1, with $\mu$, is dropped, and the particles are only allowed to be displaced, not inserted or removed. We identify the phase coexistence region, as shown in the representative phase diagram in Supplementary Fig. S1B. Here, the red line represents the binodal curve, outside which only a homogeneous phase is thermodynamically stable[67]. We also identify the critical point for the bulk coupling parameter as $J_{b,crit} \simeq 0.18\ k_BT$. This is also corroborated by the peak in the specific heat (Supplementary Fig. S1C), which diverges at $J_{b,crit}$ in the thermodynamic limit. We further verify that our simulations reproduce the expected droplet coarsening dynamics. For the average droplet size, we observe a scaling of $<R(t) \sim t^{1/3}>$ (Supplementary Fig. S1D), in agreement with the Lifshitz-Slyozov-Wagner scaling, which describes droplet coalescence via Ostwald ripening[70,71]. Together, these results confirm that our model correctly captures both the equilibrium and kinetic characteristics of classical phase separation in solution.

While we work in the canonical ensemble to identify the coexistence region and critical coupling constant, this ensemble poses a limitation in the presence of the microtubule. Condensation of tau molecules onto the surface depletes the solution, artificially limiting condensate growth, even if it is thermodynamically favorable to do so. One possible workaround is to use a sufficiently large simulation box so that the concentration in the solution is not altered significantly by the loss of particles to the surface condensate. An alternative approach is to work in a grand canonical ($\mu VT$) ensemble, which allows the number of particles to fluctuate, keeping the average concentration in the solution fixed, even as the surface condensates form and grow. We adopt the grand canonical ensemble in our simulations. Now, instead of specifying the volume fraction $\phi$ for the bulk, we specify the bulk chemical potential $\mu$ (Eq.1). First, we aim to identify the range of $\mu$ corresponding to the undersaturated regime. The phase diagram in the $J_b$- $\mu$ space is shown in Supplementary Fig. S1E, where the red line represents the first-order transition line between the dense and dilute phases, terminating at the critical point $J_{b,crit}$. This first-order transition line follows the relation $\mu = -qJ_b/2$, where $q$ is the coordination number and $J_b > J_{b,\, crit}$. Setting $\mu$ to this value is equivalent to setting the external field to zero in the standard Ising model. At values of $\mu < -qJ_b/2$, the system will be in the undersaturated regime, which is the parameter regime of our interest. Additionally, in the dilute limit, we can relate the bulk chemical potential to concentrations in the experiments via the relation

$$\mu = \mu_0 + \alpha log\left(\frac{c}{c_0}\right), \tag{2}$$

where $c_0$ is the reference concentration at which $\mu = \mu_0$ and α is a scaling parameter with units $k_BT$ [44].

**1.4. Excess density and the fraction of microtubule length covered captures the extent of tau condensation**

As described in the Main text, even in the undersaturated regime, a dense (liquid) phase can form, which is stable only in the vicinity of the surface. The extent of surface condensation of tau can be quantified using 1) the surface excess density[72], and 2) the fraction of the microtubule length covered by the condensates. The excess density is defined as

$$c_s = \frac{1}{N}\sum_{n=1}^{N}(\phi_n - \phi_b) \tag{3}$$

where $\phi_n$ is the average occupancy of the $n^{th}$ layer and $\phi_b$ is the density in the solution far away from the surface. The layers are defined radially outward from the microtubule surface, such that $n$=1 corresponds to the first layer where the molecules are directly bound to the microtubule, $n$=2 is the next outward layer and so on.

To obtain the density profile along the microtubule, we take the average density in a region of size 10x10 around the microtubule at each position along its length, and subtract the average bulk density from it. This is analogous to measuring the intensity of tau on the microtubule surface from the microscopy data, and allows comparison to the experimental results. The fraction of microtubule length covered with tau condensates is calculated as the fraction of sites having a density greater than 1.5 times the bulk density (although this threshold is arbitrary, varying it does not change the results qualitatively). The kymographs from the simulation were similarly obtained, measuring the density profile along the microtubule length as a function of time (Main text Fig. 2C). To study the disassembly kinetics of the tau condensates (Main text Fig. 2C), we first let the condensate form and equilibrate with some value of bulk chemical potential $\mu$. Then, using this as our initial configuration, we evolve the system with a much lower value of $\mu$, emulating the washout process observed in the experiments (see the respective figure captions for the exact parameter values).

##### 1.5. Parameter selection

In this section, we motivate our choice of parameters for the model. The average length of the microtubule in the experiments is about 10$\mu m$. The size of a tubulin dimer is ~8 nm[73], with a stoichiometry between 1:2 and 1:5 tau: tubulin[55], giving us around 250-625 binding sites for tau along each protofilament. We take the microtubule to have $M$=300 binding sites along each protofilament.

Since the tau condensates are localized at the microtubules, we can take a smaller size for the simulation box in the directions perpendicular to the microtubule axis. This is done to minimize the computation in the solution. We take $L$=26 sites. The value of the bulk interaction parameter $J_b$ is taken to be 0.2 $k_BT$ as (above the critical value of 0.18$k_BT$). This sets the range for the bulk chemical potential as $\mu < -qJ_b/2 = -2.6\ k_BT$, below which the system is undersaturated. The surface coupling parameter $J_s$ is taken to be higher than $J_b$, indicating a positive binding cooperativity. Drawing comparisons with the experiments requires us to relate the chemical potential to the concentration of tau used. The parameters for this conversion ($\mu_0$, $\alpha$ and $c_0$ in Eq.2) were obtained as fitting parameters for the tau density and coverage data in Supplementary Fig. S6B, C. The simulation time step $\Delta t$ was set to match the timescales of tau condensate assembly and disassembly in the experiments.

| Parameters | Range/Value |
|---|---|
| Bulk coupling parameter, $J_b$ | [0.18, 0.25] |
| Surface coupling parameter, $J_s$ | [0.3, 1.0] |
| Surface heterogeneity mean, $\bar{h}$ | [-0.5, 1.0] |
| Surface heterogeneity std. Deviation, $\sigma$ | [0.0, 0.5] |
| Bulk chemical potential, $\mu$ | [-4.0, -2.5] |
| Reference chemical potential, $\mu_0$ | -2.904 |
| Chemical potential scaling factor, $\alpha$ | 0.05536 |
| Reference concentration, $c_0$ *(nM)* | 4.715 |
| Number of tau binding sites along MT, $M$ | [120, 300] |
| Simulation box size perpendicular to MT, $L$ | 26 |
| Simulation time step per sweep, $\Delta t$ *(s)* | 0.035 |

Table 1. Parameter values used (units are $k_BT$ unless mentioned).

#### 2. Tau forms surface condensates on microtubules

##### 2.1. Homogeneous model

We first consider a homogeneous microtubule surface, where the binding affinity $h_i$ has the same value at all the sites. As mentioned above, in our model, condensation in solution occurs at $\mu > -2.6\ k_BT$. In the undersaturated regime, at low concentrations ($\mu \sim -2.9\ k_BT$) dense patches, referred to as condensates, form on the microtubule surface (Supplementary Fig. S2A). Following the kinetics of tau condensation on microtubules, we find that the average length of tau condensates increases linearly with time, $\langle l(t) \sim t \rangle$ (Supplementary Fig. S2B) as expected for condensate growth on a one-dimensional surface[74]. However, the condensates are not spatially pinned, as seen in the experiments.

At equilibrium, tau predominantly exhibits two states: 1) at low concentrations, it is adsorbed directly onto the microtubule as individual molecules, and 2) at higher concentrations, it forms multilayer condensates (Supplementary Fig. S2C). As the tau concentration is increased, the switch from a thin, adsorbed layer to a thick, condensed layer is sharp (Supplementary Fig. S2D). Moreover, the two states can coexist near the transition concentration, as shown in Main text Fig. 1J. This rapid switch is akin to a prewetting transition, which was first predicted by Cahn[20,22,75]. The prewetting transition is a first-order transition, where the surface state goes from a thin adsorbed layer to a thick layer in a discontinuous manner (Supplementary Fig. S2E). This is captured by a sharp jump in the excess density as we increase the concentration (bulk chemical potential $\mu$) or the surface binding affinity $h$.

We then systematically vary individual model parameters to explore their effect on the equilibrium behaviour. Varying the surface coupling term $J_s$ and the binding affinity $h$, we identify the coexistence line along which the thin and thick layers coexist (Supplementary Fig. S2G, H). Similar to the bulk, there exists a critical point at which this first-order transition line terminates, beyond which the transition from an adsorbed layer to a condensed layer is no longer sharp. This can be seen for lower values of $J_s$ (or equivalently for higher temperatures).

Next, we then vary the bulk chemical potential $\mu$ and the surface binding affinity $h$. It can be seen from the Hamiltonian (and from Supplementary Fig. S2I, J) that the coexistence line can be approximated as $\mu + h = constant$. This means that one can affect an increase in the degree of surface condensation by either making the surface more attractive (increasing $h$) or by increasing the concentration of molecules in the solution (increasing $\mu$). For a given condensation profile, the choice of $\mu$ and $h$ is not unique. We use this to set $h = 0\ k_BT$ in our model. Nevertheless, to describe the entire system uniquely (both bulk and surface states), we must specify both parameters independently.

#### 2.2. Heterogeneous model

As mentioned in the Main text, the homogeneous model fails to capture the following experimental observations: 1) the condensates form in the same place for a given microtubule, 2) the condensates stay pinned to their location. To explain these observations, we introduce heterogeneity in the binding affinity profile. The affinity at each position along the microtubule $i$ is drawn randomly from a normal distribution $h_i \sim Normal\ (\bar{h},\ \sigma)$. As a simplification, all the 12 sites around the microtubule for a given position $i$ have the same binding affinity. As described in the previous section, we set $\bar{h} = 0$, and choose $\sigma = 0.2\ k_BT$ (these parameters are chosen to capture the behaviour shown by taxol-stabilized microtubules; parameters for GDP- and GMPCPP-stabilized microtubules are given in the following section).

Similar to the homogeneous model, tau is adsorbed onto the microtubules at lower concentrations and forms multilayered condensates at higher concentrations. However, the transition from thin to thick layer is no longer sharp, and the system exhibits coexistence between the two states for a wider range of concentrations (Supplementary Fig. S3B-C, E-F). Importantly, the tau condensates remain pinned to their location on the microtubule, similar to the experiments (Supplementary Fig. S3A). We next examine how the mean binding affinity $\bar{h}$ and the degree of heterogeneity $\sigma$ affect the kinetic and the equilibrium behaviour. As expected, increasing $\bar{h}$ leads to an increase in tau condensation on the microtubules. Increasing the heterogeneity $\sigma$ broadens the region of coexistence between the thin and thick layers. This can be seen in Supplementary Fig. S3D, as the transition is sharper for lower values of $\sigma$, with $\sigma = 0\ k_BT$ (homogeneous surface) showing a very sharp transition. Furthermore, increasing $\sigma$ results in the shorter condensate lengths on average (Supplementary Fig.S3D).

#### 2.3. Heterogeneous model captures experimental observations with different kinds of microtubules

In this section, we further compare the heterogeneous model to the experimental observations. GDP-microtubules have a more compacted lattice, and GMPCPP-stabilized microtubules have an irreversibly extended lattice[6]. This results in different values of tau binding affinities as well as different values of tau cooperativity. We can incorporate this into our model by varying both $h$ and $J_s$. Here, we take the value of $J_s$ to be the same across all three microtubule types, and consider different values of $\bar{h}$. As mentioned in the previous section, we chose $\bar{h} = 0.0\ k_BT$ for taxol-stabilized microtubules. For GDP-microtubules, we set $\bar{h} = 0.3\ k_BT$, and for GMPCPP-stabilized microtubules we set $\bar{h}$ = -0.1 $k_BT$. The level of heterogeneity, governed by $\sigma$, is taken to be the same (0.2 $k_BT$) for all three microtubules. As mentioned in the Main text, tau covers the entire GDP-microtubule surface even at very low concentrations, whereas GMPCPP-stabilized microtubules show a dense, uniform layer only at very high concentrations (Main text Fig. 1B). The heterogeneous model captures the equilibrium and kinetic behaviour across different concentrations for all three microtubule types very well (Supplementary Fig. S4).

#### 3. Previously proposed 1D model captures features of tau condensation on microtubules

A recent study investigated the interaction between the pioneer transcription factor KlF4 and lambda-DNA[44]. Similar to tau on taxol-stabilized microtubules, KlF4 showed a bimodal distribution of intensities on the DNA. The two states of KlF4 were identified as an adsorbed state and a condensed state, with the switch between them attributed to a prewetting transition. The resulting condensation patterns were shown to be correlated to the DNA sequence and were explained using a one-dimensional model. In this model, each site along the 1D lattice is assumed be in one of two states: adsorbed or condensed. Here, we show that the same model can be used to capture the kinetic and equilibrium properties of tau condensation on microtubules as well.

In this model, the microtubule is considered as a one-dimensional lattice of $M$ sites, where each site corresponds to the local wetting state of tau: -1 for a thin adsorbed layer and +1 for a thick condensed layer (Supplementary Fig. S8A). This model is equivalent to a one-dimensional random-field Ising model[76]. The effective free energy of the system is given by

$$E = -J \sum_{\langle i,j \rangle} s_i \, s_j - \sum_i (h + h_i) \, s_i \tag{4}$$

where the index $i$=1,2,...,$M$ and $<i,j>$ indicate the nearest neighbouring pairs. The nearest-neighbour interaction is governed by the parameter $J$, and it reflects the interfacial tension in the wetted layers (related to $J_b$ and $J_s$ in our 3D model). Each site has some affinity to form a thin or thick layer, represented by the term $h_i$. The parameter $h$ relates to the bulk chemical potential of the proteins, which in the dilute limit can be given as $h = h_0 + \alpha \log(C/C_0)$. Here $C$ is the protein concentration in solution, and $C_0$ is a reference concentration. $h_0$ is the chemical potential at concentration $C_0$ and $\alpha$ is a proportionality constant, having the same dimensions as $h$. For simplification, we set $h_0 = 0$, and $C_0$ is the concentration at which roughly half of the microtubule is covered by the condensed region.

As in the 3D lattice gas model, we consider two kinds of surfaces: a homogeneous surface with a constant binding affinity, and a heterogeneous surface with noisy binding affinities, sampled from a Normal distribution. For the case of homogeneous binding affinities, we set $h_i$=0 and use the transfer matrix method[77] to obtain an expression for the fraction of microtubule covered by the tau condensate[44] as

$$\phi = \frac{1}{2}\left(1 - \frac{1+\left(\frac{C}{C_0}\right)^{-2\beta\alpha}}{\sqrt{1+4e^{(-4\beta J)}+\left(\frac{C}{C_0}\right)^{-4\beta\alpha}-2\left(\frac{C}{C_0}\right)^{-2\beta\alpha}}}\right) \quad (5)$$

For a heterogeneous surface, we consider the site-dependent affinities $h_i$ to be drawn randomly from a normal distribution with zero mean and standard deviation σ. We numerically compute the microtubule fraction in the condensed state, using $J$, $\alpha$ and $C_0$ as fitting parameters (see figure captions for parameter values). To study the temporal behaviour of the system, we employ the kinetic Monte Carlo method. The rate of switching any given site using the energy given in Eq.4 and the detailed balance condition can be defined as in[44]. The rate of switching the $j^{th}$ site from +1 to -1 is taken as $\omega_j = Ae^{-(2\beta(\delta+1)H_j)}$, and for switching from -1 to +1 as $\omega_j = Ae^{-(2\beta\delta H_j)}$, where $H_j$ is the local field at site $j$, and is given by $H_j = -J(s_{j-1} + s_{j+1}) - (h + h_i)$. The parameters $A$ and δ dictate the timescales of condensate formation and growth, and are estimated by comparing with the experimental timescales.

We find that a homogeneous surface can explain the equilibrium behaviour for the surface fraction as a function of tau concentration (Supplementary Fig. S8B). However, since the binding affinity is the same everywhere on the surface, the homogeneous surface cannot give rise to stable pinned condensates (Supplementary Fig. S8C). Additionally, across independent simulation trials, the homogeneous surface does not nucleate tau condensates in the same location, as is observed in the experiments (Main text). However, in contrast, a heterogeneous surface can capture both the equilibrium and kinetic behaviour (Supplementary Fig. S8D).

While these results are in agreement with those obtained from the three-dimensional lattice-gas model, the latter is more general. The one-dimensional model assumes that there are two states: adsorbed and condensed, and the switch between them is a sharp transition, whereas in our lattice gas model, these states emerge due to tau-tau and tau-microtubule interactions.

#### Supplementary information references


65. Gustke, N., Trinczek, B., Biernat, J., Mandelkow, E.-M. & Mandelkow, E. Domains of tau protein and interactions with microtubules. *Biochemistry* **33**, 9511–9522 (1994).

66. Drechsel, D. N., Hyman, A. A., Cobb, M. H. & Kirschner, M. W. Modulation of the dynamic instability of tubulin assembly by the microtubule-associated protein tau. *Mol. Biol. Cell* **3**, 1141–1154 (1992).

67. Flory, P. J. Thermodynamics of high polymer solutions. *J. Chem. Phys.* **10**, 51–61 (1942).

68. Binder, K. & Landau, D. P. Monte Carlo simulation of wetting transitions in the ferromagnetic Ising model. *J. Appl. Phys.* **57**, 3306–3308 (1985).

69. Binder, K., Landau, D. P. & Wansleben, S. Wetting transitions near the bulk critical point: Monte Carlo simulations for the Ising model. *Phys. Rev. B* **40**, 6971–6979 (1989).

70. Lifshitz, I. M. & Slyozov, V. V. The kinetics of precipitation from supersaturated solid solutions. *J. Phys. Chem. Solids* **19**, 35–50 (1961).

71. Wagner, C. Theorie der alterung von niederschlägen durch umlösen (ostwald-reifung). *Z. Für Elektrochem. Berichte Bunsenges. Für Phys. Chem.* **65**, 581–591 (1961).

72. Binder, K. & Landau, D. P. Wetting and layering in the nearest-neighbor simple-cubic Ising lattice: A Monte Carlo investigation. *Phys. Rev. B* **37**, 1745–1765 (1988).

73. Desai, A. & Mitchison, T. J. Microtubule polymerization dynamics. *Annual Review of Cell and Developmental Biology* vol. 13 83–117 (1997).

74. Pombo-García, K., Adame-Arana, O., Martin-Lemaitre, C., Jülicher, F. & Honigmann, A. Membrane prewetting by condensates promotes tight-junction belt formation. *Nature* **632**, 647–655 (2024).

75. Ebner, C. & Saam, W. F. New Phase-Transition Phenomena in Thin Argon Films. *Phys. Rev. Lett.* **38**, 1486–1489 (1977).

76. Blossey, R., Kinoshita, T. & Dupont-Roc, J. Random-field Ising model for the hysteresis of the prewetting transition on a disordered substrate. *Phys. Stat. Mech. Its Appl.* **248**, 247–272 (1998).

77. Chaikin, P. M. & Lubensky, T. C. *Principles of Condensed Matter Physics*. (Cambridge University Press, Cambridge, 1995).

#### Supplementary figures

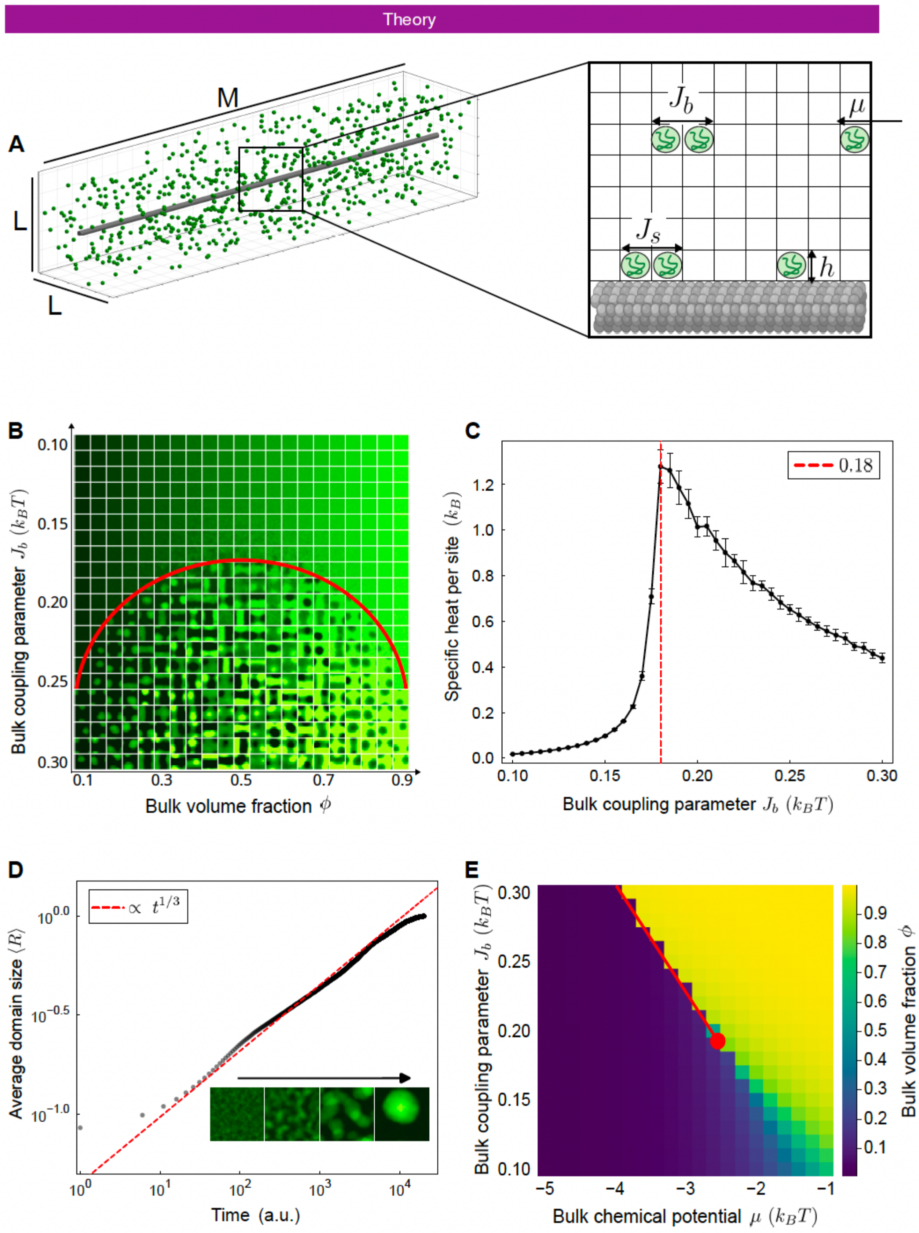


***Supplementary Fig. S1 - Characterizing phase separation in the bulk.***

***(A)*** *Schematic of the lattice-gas model of tau interacting with microtubules.* ***(B)*** *Representation of the phase diagram in an NVT ensemble. The red line represents the binodal, inside which the dense and dilute phases coexist at equilibrium.* ***(C)*** *Specific heat as a function of the bulk interaction parameter* $J_b$*.* ***(D)*** *Kinetics of the droplet growth is captured in the model (inset). This agrees with the Lifshitz-Sylozov scaling relation (averaged over 30 simulations,* $J_b = 0.3\ k_BT$*,* $\phi=0.2$*).* ***(E)*** *The bulk coexistence line in a* $\mu VT$ *ensemble. The red line represents the threshold at which the dense and dilute phases can coexist, given by* $\mu = -qJ_b/2$*. This line terminates in the critical point* $J_{b,crit}$*. (Simulations shown here were carried out for box size 40x40x40).*

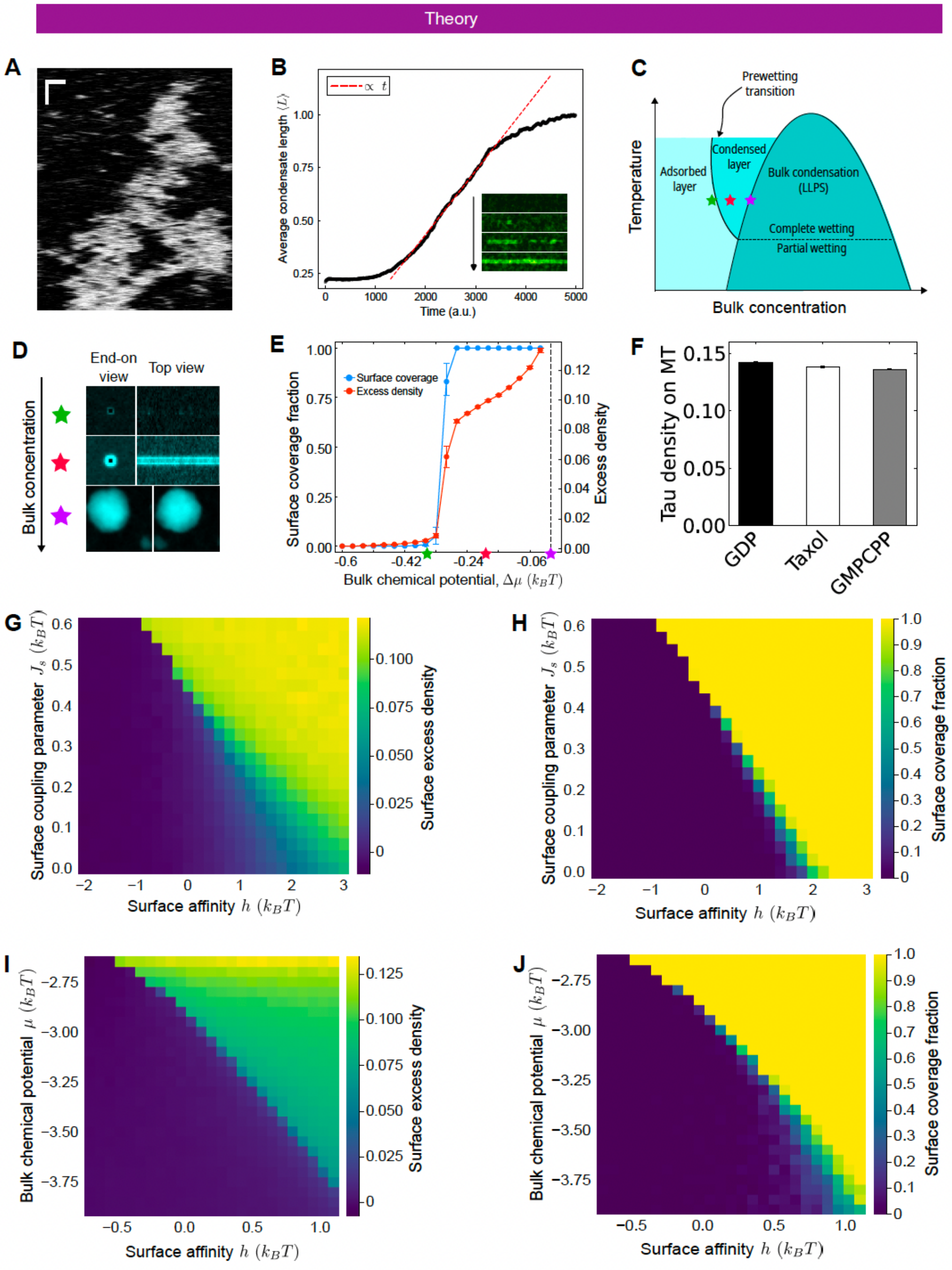


***Supplementary Fig. S2 - Homogeneous model.***

***(A)*** *Kymograph showing tau condensation on a homogeneous microtubule. The condensates are not pinned to their location. (horizontal scale bar is 1μm and vertical is 5s)* ***(B)*** *The kinetics of the condensate growing on the microtubule show the size increasing linearly with time.* ***(C)*** *Schematic of the phase diagram showing the wetting and prewetting transition.* ***(D)*** *Green star: adsorbed tau ($\mu$ = -2.92 $k_BT$), red star: condensed tau ($\mu$ = -2.8 $k_BT$) and purple star: bulk droplet ($\mu$ = -2.6 $k_BT$). (E) The excess density (orange) and the* *fraction of microtubule surface covered by condensate (blue) show a sharp jump with increasing bulk chemical potential. $\Delta\mu = 0$ corresponds to bulk phase separation, indicated by a dashed line. The stars represent the concentrations mentioned in C.* ***(F)*** *Simulation results showing tau intensities at the microtubule for three different surfaces, representing GDP-, taxol-stabilized and GMPCPP-stabilized microtubules. ($h_i = 0.3\ k_BT$, $h_i = 0.0\ k_BT$, $h_i = -0.1\ k_BT$ respectively).* ***(G-H)*** *The excess density and the fraction of surface covered, respectively, vary with the surface coupling parameter $J_s$ and the surface affinity h. Similar to the bulk, there exists a first-order transition on the surface, where the surface densities show a sharp jump upon varying the affinity. This is a characteristic of a prewetting transition ($\mu = -2.9\ k_BT$).* ***(I-J)*** *The excess density and the fraction of surface covered, respectively, as a function of the bulk chemical potential $\mu$ and the surface affinity h. The sharp jump in the surface densities as a function of $\mu$ is another characteristic of the prewetting transition. A slice along the line at h = 0.0 is shown in (E). Wherever not explicitly specified, the simulations were performed with $J_b = 0.2\ k_BT$, $J_s = 0.5$ kBT, hi = 0.0 $k_BT$ (representing taxol microtubules).*

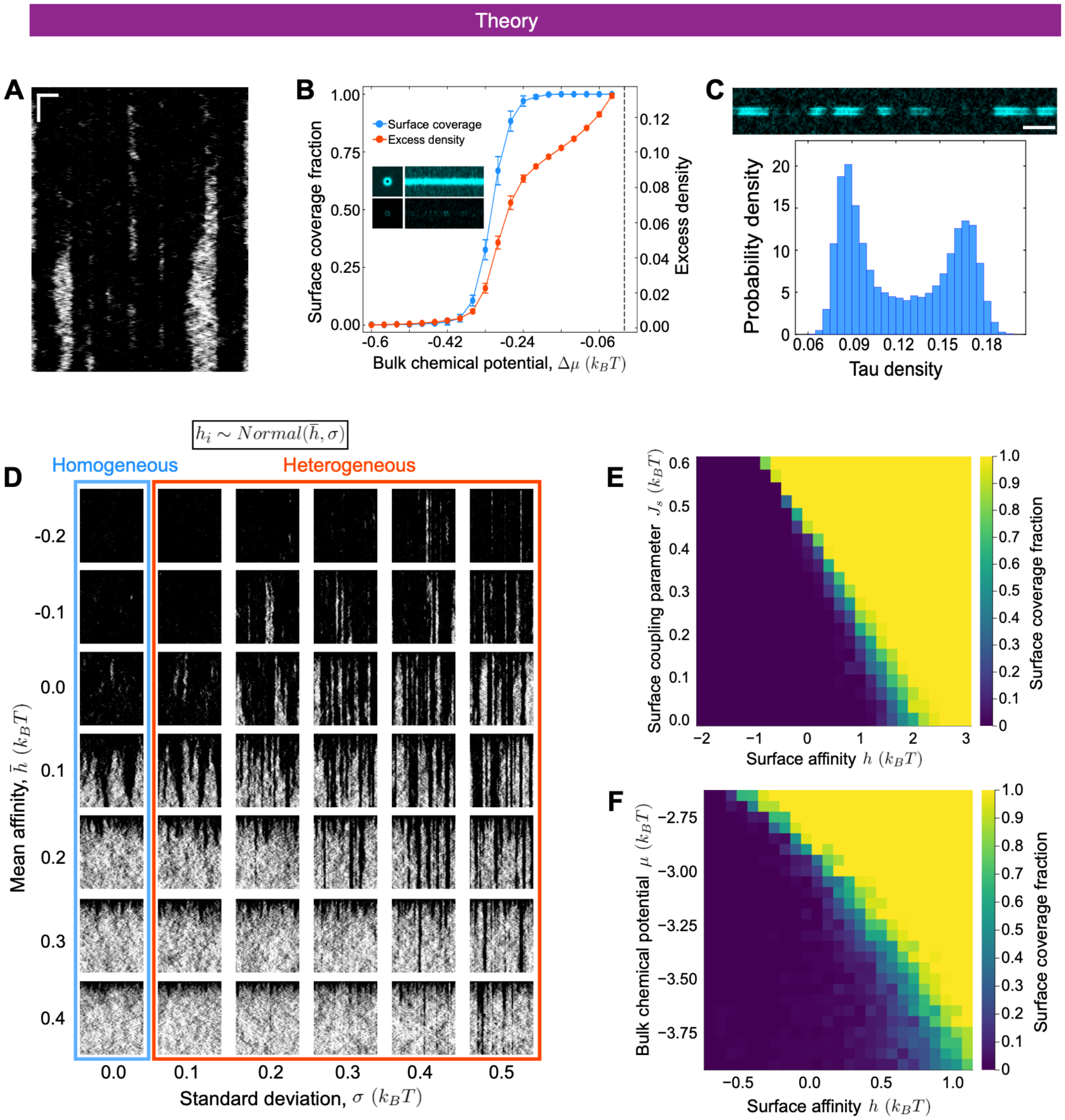


***Supplementary Fig. S3 - Heterogeneous model.***

**(A)** *Kymograph showing tau condensation on a heterogeneous microtubule. In contrast to the homogeneous model, here the condensate is spatially pinned (horizontal scale bar is 1μm and vertical is 5s).* ***(B)*** *Corresponding plot to Fig. S2E, showing the excess density (orange) and the fraction of microtubule surface covered by condensate (blue). Introducing heterogeneity destroys the sharp jump observed with a homogeneous surface.* ***(C)*** *The heterogeneous model also shows the coexistence of the adsorbed and condensed states. Intensity distribution shows a bimodal shape (obtained over 100 simulations) (μ = -2.925 $k_BT$).* ***(D)*** *Effect of parameters $\bar{h}$ and σ. The first column, with σ=0.0, corresponds to a homogeneous microtubule. (total time for each microtubule is 500s, and the length is 6μm).* ***(E-F)*** *Effect of introducing heterogeneities as seen in the equilibrium states. The coexistence region gets broader due to the introduction of heterogeneities in the binding affinity (μ = -2.723 $k_BT$ in E, and $J_s$ = 0.5 $k_BT$ in F). Simulations were performed with $J_b$ = 0.2 $k_BT$, $J_s$ = 0.5 $k_BT$, and $h_i$ ~ N (0.0, 0.2) $k_BT$ unless explicitly specified.*

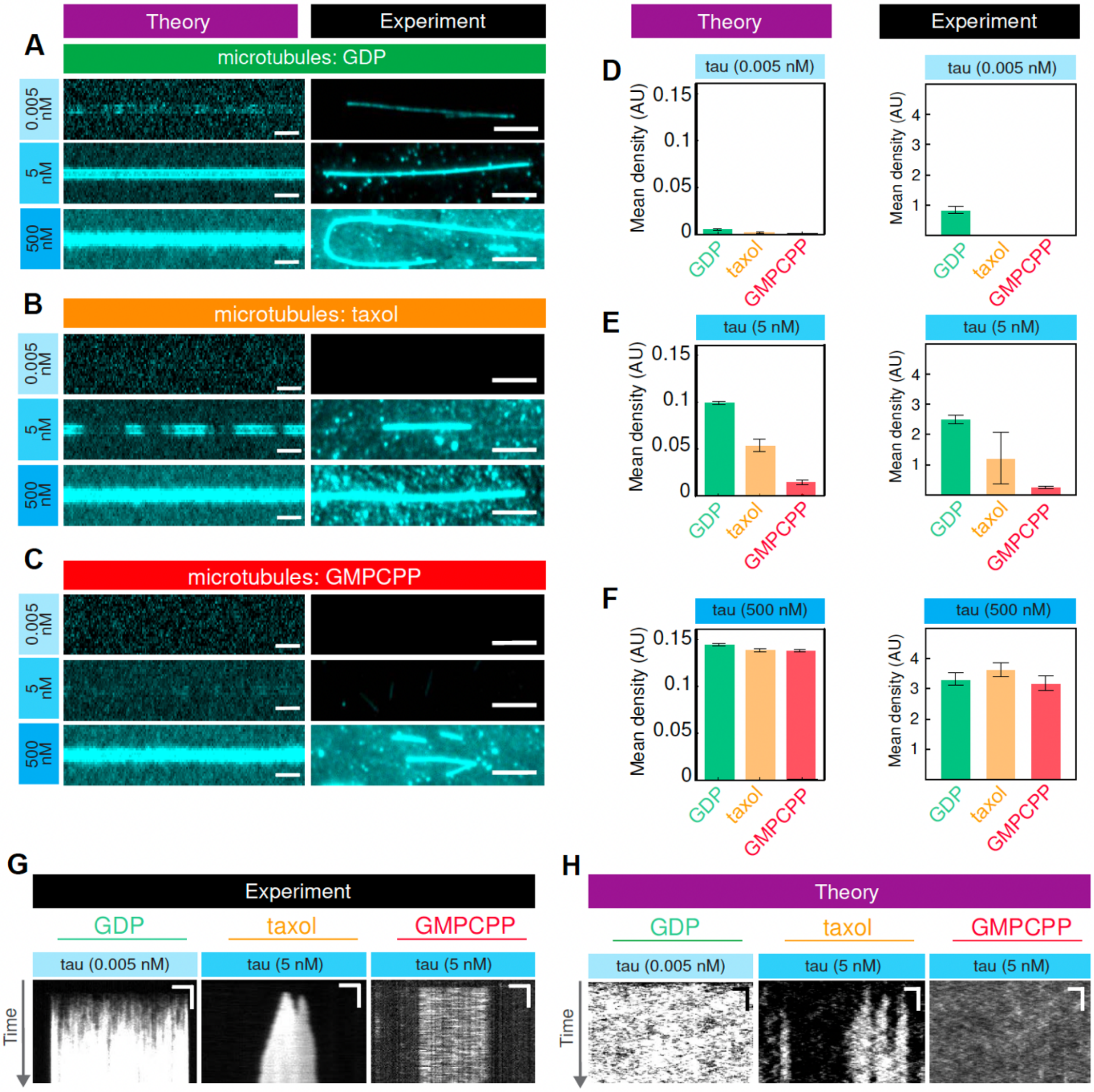

Theory
Experiment
A
microtubules: GDP
0.005 nM
5 nM
500 nM
B
microtubules: taxol
C
microtubules: GMPCPP
D
tau (0.005 nM)
E
tau (5 nM)
F
tau (500 nM)
Mean density (AU)
GDP
taxol
GMPCPP
G
Experiment
H
Theory
Time

***Supplementary Fig. S4 - Heterogeneous model captures the equilibrium and kinetic behaviour with various microtubules.***

***(A-C)*** *Comparison between simulation results and experimental snapshots of the equilibrium state for GDP- ($\bar{h}$ = 0.3 $k_BT$), taxol- ($\bar{h}$ = 0.0 $k_BT$), and GMPCPP-stabilized microtubules ($\bar{h}$ = -0.1 $k_BT$) for 0.005nM, 5nM or 500nM tau. Scale bars, experiment: 5 µm, theory: 1 µm.* ***(D-F)*** *Quantification of the mean density of tau on the microtubules for GDP-, taxol- and GMPCPP-stabilized microtubules as predicted by the model (left), and as measured in the experiments (right).* ***(G)*** *Experimental kymographs showing tau states on different microtubules (same as shown in Main text Fig.1) Scale bars: horizontal, 5 µm; vertical, 20 s* ***(H)*** *Simulation kymographs showing tau states on different microtubules. Scale bars: horizontal, 1 µm; vertical, 20 s*

*(All simulations were performed with $J_b$ = 0.2 $k_BT$, $J_s$ = 0.5 $k_BT$, $\sigma$= 0.2 $k_BT$ )*

*All data in D, E, F are presented as mean ± SD (error bars).*

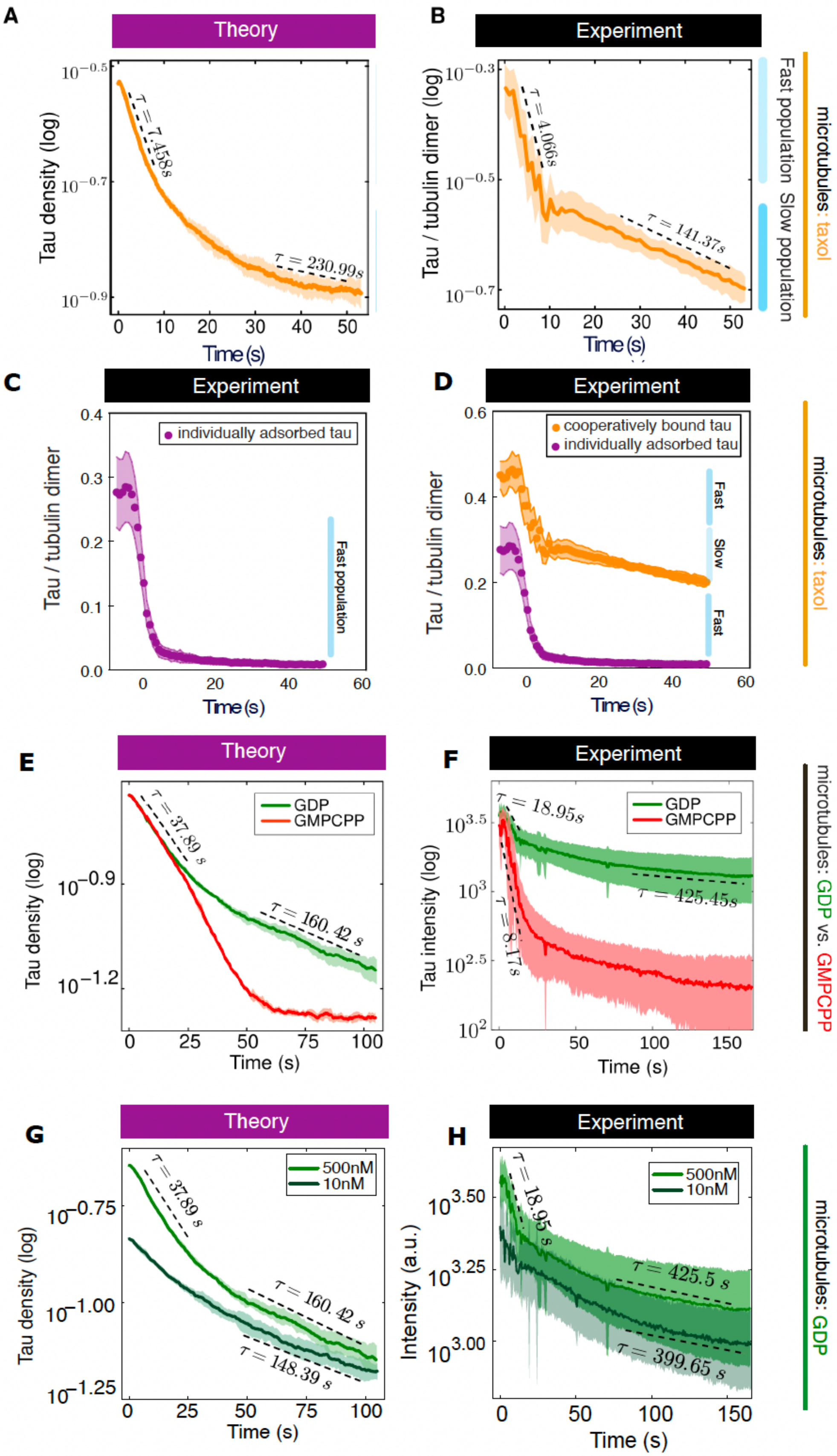
A
Theory
Tau density (log)
τ = 7.458s
τ = 230.99s
Time (s)
B
Experiment
Tau / tubulin dimer (log)
τ = 4.066s
τ = 141.37s
Time (s)
Fast population
Slow population
microtubules: taxol
C
Experiment
individually adsorbed tau
Tau / tubulin dimer
Fast population
Time (s)
D
Experiment
cooperatively bound tau
individually adsorbed tau
Tau / tubulin dimer
Fast
Slow
Fast
Time (s)
microtubules: taxol
E
Theory
GDP
GMPCPP
τ = 37.89 s
τ = 160.42 s
Tau density (log)
Time (s)
F
Experiment
GDP
GMPCPP
τ = 18.95s
τ = 8.17s
τ = 425.45s
Tau intensity (log)
Time (s)
microtubules: GDP vs. GMPCPP
G
Theory
500nM
10nM
τ = 37.89 s
τ = 160.42 s
τ = 148.39 s
Tau density (log)
Time (s)
H
Experiment
500nM
10nM
τ = 18.95 s
τ = 425.5 s
τ = 399.65 s
Intensity (a.u.)
Time (s)
microtubules: GDP

***Supplementary Fig. S5 - Condensate disassembly reveals two kinetically distinct populations of tau***

***(A)*** *Simulation of tau density during disassembly, revealing fast- and slow-turnover populations identified by a double-exponential fit.* ***(B)*** *Experimental data showing time-traces of tau-meGFP density during washout, showing fast- and slow-turnover components within condensates (160 nM tau). N=9 condensate regions.* ***(C)*** *Experimental time traces of tau–meGFP density during washout, revealing only the fast-turnover component outside the tau condensate region (tau concentration, 160 nM).* ***(D)*** *Comparison of tau–meGFP washout kinetics inside (orange) and outside (purple) the tau condensates. The condensate region exhibits both fast- and slow-turnover components, whereas only the fast-turnover component is observed outside the condensate region (tau concentration, 160 nM). N=9 condensate regions and N=5 non-condensate regions.* ***(E)*** *Model prediction for condensate disassembly when comparing microtubules with different surface properties, where the condensate is allowed to first form with 500 nM tau concentration. GDP-microtubules are assigned to have average affinity* $\bar{h} = 0.3\ k_BT$ *and for GMPCPP-stabilized microtubules* $\bar{h} = -0.1\ k_BT$*. Model predicts that the fast timescale should be similar in both the cases.* ***(F)*** *Experimental measurements of tau (500 nM) disassembly kinetics on GDP- and GMPCPP-stabilized microtubules, showing similar timescales for the fast population.* ***(G)*** *Model prediction for condensate disassembly when comparing two different tau concentrations (500 and 10 nM tau) on GDP microtubule. All simulations were performed with* $J_b = 0.2\ k_BT$, $J_s = 0.5\ k_BT$, $\sigma = 0.2\ k_BT$ *.* ***(H)*** *Experimental measurements of tau (500 and 10 nM) disassembly kinetics on GDP microtubules.*

*All data in A-H is presented as mean (solid line) ± SD (shaded areas).*

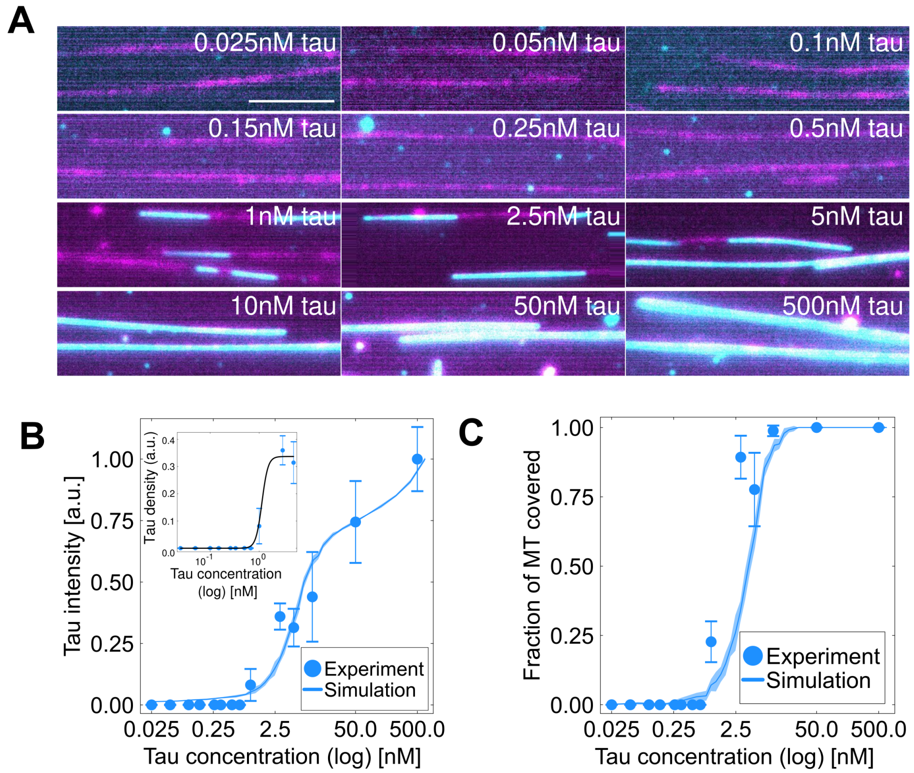


***Supplementary Fig. S6 - Tau titration on microtubules***

***(A)*** *Fluorescence snapshots for different concentrations of tau on taxol-stabilized microtubules.* ***(B)*** *Theory and experimental comparison of the tau density with increasing tau concentration. Inset shows fit using a Hill function ($Ac^n/(k^n+c^n)$), with A = 0.337, n=7 and k = 1.137 nM).* ***(C)*** *Theory and experiment comparison showing the fraction of microtubule length covered by the tau condensates. We fit the model parameters $\mu_0$, $\alpha$ and $c_0$, in Eq. 2, to convert the chemical potential $\mu$ to molecular concentrations using the experimental data in B and C. The fitting parameters obtained for this conversion are $\mu_0 = -2.904\ k_BT$, $\alpha = 0.05536\ k_BT$, and $c_0 = 4.715$ nM.*

*All data in B, C is presented as mean (dot) ± SD (error bars).*

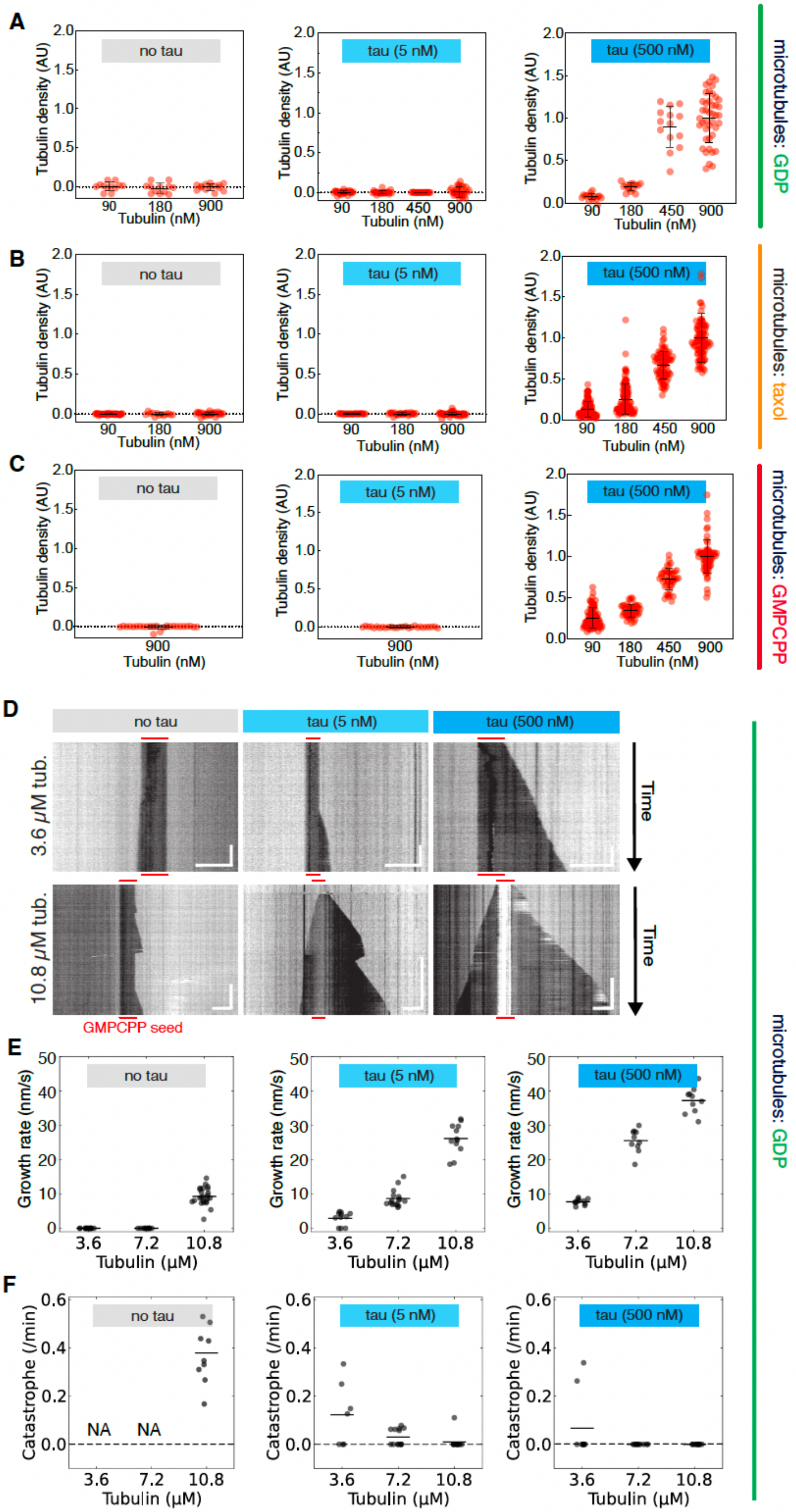
A
B
C
D
E
F
no tau
tau (5 nM)
tau (500 nM)
Tubulin density (AU)
Tubulin (nM)
microtubules: GDP
microtubules: taxol
microtubules: GMPCPP
3.6 μM tub.
10.8 μM tub.
Time
GMPCPP seed
Growth rate (nm/s)
Tubulin (μM)
Catastrophe (/min)
NA
NA

***Supplementary Fig. S7 - Tubulin partitions into microtubule-condensed tau and modulates microtubule dynamics.***

***(A)*** *Quantification of free tubulin fluorescence intensity on GDP-microtubules at varying tau (0, 5, and 500 nM) and tubulin (90, 180, 450, and 900 nM) concentrations. N=15–40 microtubules per condition across 3 independent experiments.* ***(B)*** *Quantification of free tubulin fluorescence intensity on taxol-stabilized microtubules at varying tau (0, 5, and 500 nM) and tubulin (90, 180, 450, and 900 nM) concentrations. N=25–80 microtubules per condition across 3 independent experiments.* ***(C)*** *Quantification of free tubulin fluorescence intensity on GMPCPP-stabilized microtubules at varying tau (0, 5, and 500 nM) and tubulin (90, 180, 450, and 900 nM) concentrations. N=25–60 microtubules per condition across 3 independent experiments.* ***(D)*** *Kymographs showing microtubule growth from GMPCPP seeds following addition of 3.6 or 10.8 µM tubulin in the presence of 0, 5, and 500 nM tau–mNG. Scale bars: 5 µm (horizontal), 3 min (vertical).* ***(E)*** *Microtubule growth rates for conditions in (D). N=12–24 microtubules per condition across 2 independent experiments.* ***(F)*** *Catastrophe rates for conditions in (D). N=10–15 microtubules per condition across 2 independent experiments.*

*All data in A, B, C, E, F is presented as mean (line) ± SD (error bars).*

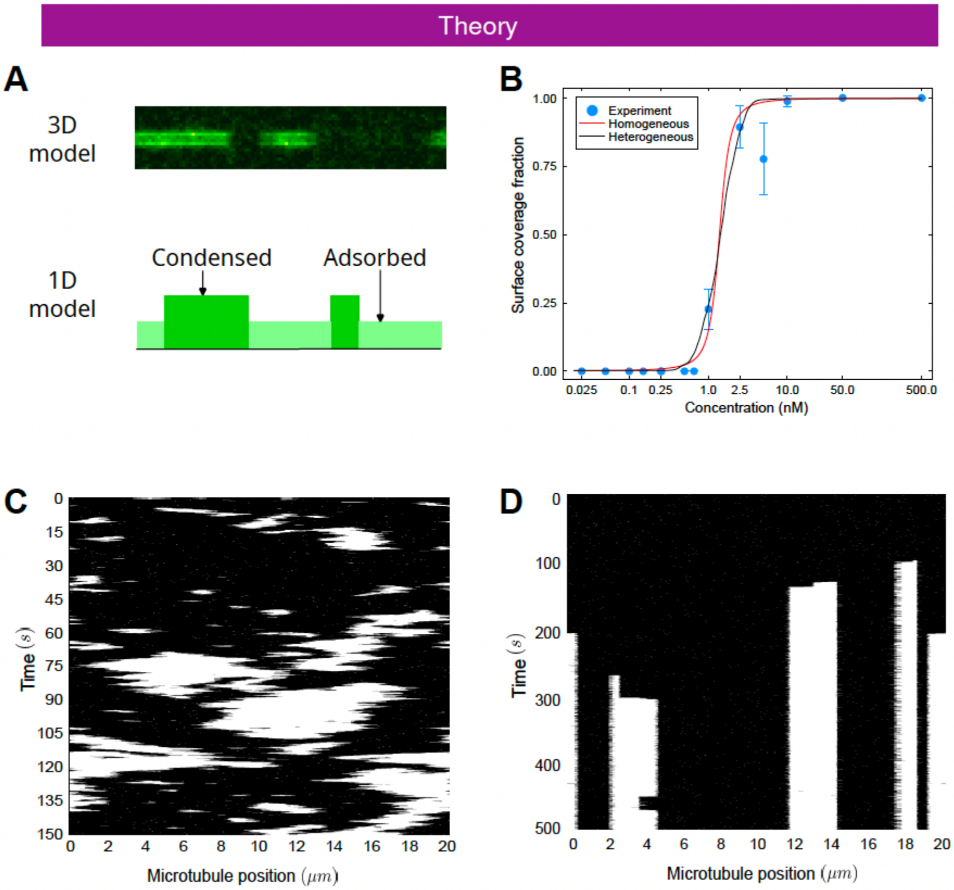


***Supplementary Fig. S8 - Previously proposed one-dimensional model for surface condensation.***
***(A)*** *Coarse-graining the 3D model gives us the 1D model of condensation of tau proteins on microtubules.* ***(B)*** *Both homogeneous and heterogeneous models can explain the equilibrium data (homogeneous model, red: $J = 2.3\ k_BT$, $\alpha = 0.031\ k_BT$, $c_0 = 1.382$ nM; heterogeneous model, black: $J = 4.5\ k_BT$, $\alpha = 0.52\ k_BT$, $c_0 = 1.43$ nM.* ***(C)*** *The homogeneous model cannot give rise to pinned islands ($A = 100.0$, $\delta = 0.1$).* ***(D)*** *Heterogeneous model gives rise to stable, pinned islands ($h_i$ drawn from $N$ (0.0, 1.4), $A = 500.0$, $\delta = 0.001$). All data in B is presented as mean (dot) ± SD (error bars).*
